# RESOLVING BIOLOGY’S DARK MATTER: SPECIES RICHNESS, SPATIOTEMPORAL DISTRIBUTION, AND COMMUNITY COMPOSITION OF A DARK TAXON

**DOI:** 10.1101/2024.05.07.592951

**Authors:** Emily Hartop, Leshon Lee, Amrita Srivathsan, Mirkka Jones, Pablo Peña-Aguilera, Otso Ovaskainen, Tomas Roslin, Rudolf Meier

## Abstract

**Background:** Zoology’s dark matter comprises hyperdiverse, poorly known taxa that are numerically dominant but largely unstudied, even in temperate regions where charismatic taxa are well understood. It is everywhere, but high diversity, abundance, and small size have historically stymied its study. We demonstrate how entomological dark matter can be elucidated using high-throughput DNA barcoding (“megabarcoding”). We reveal the high abundance and diversity of scuttle flies (Diptera: Phoridae) in Sweden using 31,800 specimens from 37 sites across four seasonal periods. We investigate the number of scuttle fly species in Sweden and the environmental factors driving community changes across time and space.

**Results:** Swedish scuttle fly diversity is much higher than previously known, with 549 mOTUs (putative species) detected, compared to 374 previously recorded species. Hierarchical Modelling of Species Communities reveals that scuttle fly communities are highly structured by latitude and strongly driven by climatic factors. Large dissimilarities between sites and seasons are driven by turnover rather than nestedness. Climate changes are predicted to significantly affect the 47% of species that show significant responses to mean annual temperature. Results were robust whether using haplotype diversity or species-proxies (mOTUs) as response variables. Additionally, species-level models of common taxa adequately predict overall species richness.

**Conclusions:** Understanding the bulk of the diversity around us is imperative during an era of biodiversity loss. We show that dark insect taxa can be efficiently characterized and surveyed with megabarcoding. Undersampling of rare taxa and choice of operational taxonomic units do not alter the main ecological inferences, making it an opportune time to tackle zoology’s dark matter.

## Background

> *If we go on the way we have, the fault is our greed and if we are not willing to change, we will disappear from the face of the globe, to be replaced by the insect.* – Jacques Yves Cousteau

Biodiversity loss in the Anthropocene is driven by changes in climate, land use, but also species introductions (Díaz et al. 2006; Cardinale et al. 2012). Such loss can result in the concomitant decline in ecosystem services vital to society (Mace et al. 2018; Lu et al. 2020) and may ultimately disrupt global supply chains and food security (WEF, 2020). Accurate monitoring of biodiversity is therefore a global priority (Naeem et al. 2016).

A crucial first step to monitoring biodiversity is obtaining robust quantitative baseline data. Most biologists would consider this to be data on species diversity, abundance, and biomass for those taxa that contribute substantially to these quantitative metrics. However, most biodiversity studies cover only a few well-studied groups that are relatively easily identified and quantified (e.g., birds, mammals, amphibians, bees, and butterflies). Such charismatic taxa are then used as proxies for all taxa in a region (Ricklefs 2006; Wiens 2007; Svenning et al. 2008; Ollerton 2017; Garda et al. 2018; Bell et al. 2019; Bogoni et al. 2020; Orr et al. 2020) instead of basing our understanding of global biodiversity on a broad and unbiased representation of biodiversity covering a wide range of traits and responses to environmental change (Bar-On et al. 2018, Goodsell et al. 2024).

A key component of global biodiversity are “dark taxa”, i.e., taxonomic groups for which less than 10% of the diversity is described and the species diversity is estimated to be upwards of 1000 (Hartop et al. 2022). Such poorly known groups do not only inhabit inaccessible realms like the deep sea (Rabone et al. 2023) but also the terrestrial habitats in which we live. A recent study (Srivathsan et al. 2023) revealed that 20 insect families (of which 10 belong to Diptera) account for >50% of local species diversity. Alarmingly, the very same families suffered from extreme taxonomic neglect and are therefore poorly represented in biodiversity surveys. Identifying and tackling the diversity of these dark taxa with scalable techniques thus emerges as an urgent priority for biodiversity science.

Large-scale studies of dark taxa have only become feasible in recent years due to advancements in sequencing technologies coupled with efficient single-specimen DNA barcoding workflows (Srivathsan et al. 2019a; Yeo et al. 2020; Srivathsan et al. 2021; Hartop et al. 2022; Meier et al. 2024; Chua et al., 2023: “megabarcoding”). Such workflows allow large numbers of specimens to be processed and sorted to putative species (mOTUs), while providing exact specimen counts, and vouchers for subsequent morphological, taxonomic, and biological work.

Scuttle flies in the family Phoridae (Diptera) have been considered the seventh most speciose and abundant insect family globally (Srivathsan et al. 2023). However, to date only ca. 4000 species have been described, although their actual diversity may be two orders of magnitude larger (Srivathsan et al. 2019a). In addition to their extreme species richness, scuttle flies occupy a wide range of ecological niches, from herbivores and predators to scavengers, parasitoids, and parasites (reviewed by Disney 1994). Nonetheless, previous ecological studies focusing on scuttle flies (Durska 1996, 2001, 2002, 2003, 2006, 2013, 2020; Durska et al. 2005) have been of limited scope due to time consuming morphological identification methods.

In this study, we use sorting with megabarcoding to generate the data for answering fundamental questions about this dark taxon. We ask how many species of scuttle flies occur in Sweden, how are they distributed across time and space, and what environmental variables drive their distribution. To test whether the choice of species and species delimitation method will affect the results, we carry out the same analyses using both mOTUs as species-proxies and haplotype diversity, and test whether rare species influence the overall inferences. We show how that “Dark Taxon Zoology” can quickly yield the answers urgently needed in the Anthropocene.

## Results

### Diversity

We obtained a total of 31 739 *COI* barcodes belonging to 2 697 haplotypes from scuttle fly samples from 37 sites (Fig. 1, Supp. Fig. S1) and four seasonal time periods (Supp. Fig. S2) across Sweden (Supp. Table S1). At the species threshold (1.7%) we detected 549 mOTUs. Species accumulation curves revealed that scuttle fly species diversity was incompletely sampled overall and across sites, horticultural zones and time periods (Fig. 2, Supp. Figs. S3, S4). Between 38 and 145 species were observed per site, with Chao1 richness estimates per site ranging from 83 to 244 species (Fig. 2b,c). Regional richness estimates varied between nearly 400 species estimated in the southernmost coastal zone (Zone 1), to fewer than 200 estimated for the alpine zone (Supp. Fig. S3). Midsummer and late-summer time-periods were both characterised by richness estimates of around 450 species, while the late spring and offseason time-periods showed lower species richness estimates of around 300 species (Supp. Fig. S4). Total species richness in Sweden was estimated at between 652 species (for Chao1, Fig. 2a) and 713 species (combined non-parametic estimator (CNE) in Ronquist et al. 2020), suggesting that 100-160 species of scuttle flies remain to be sampled.

**Figure 1.**
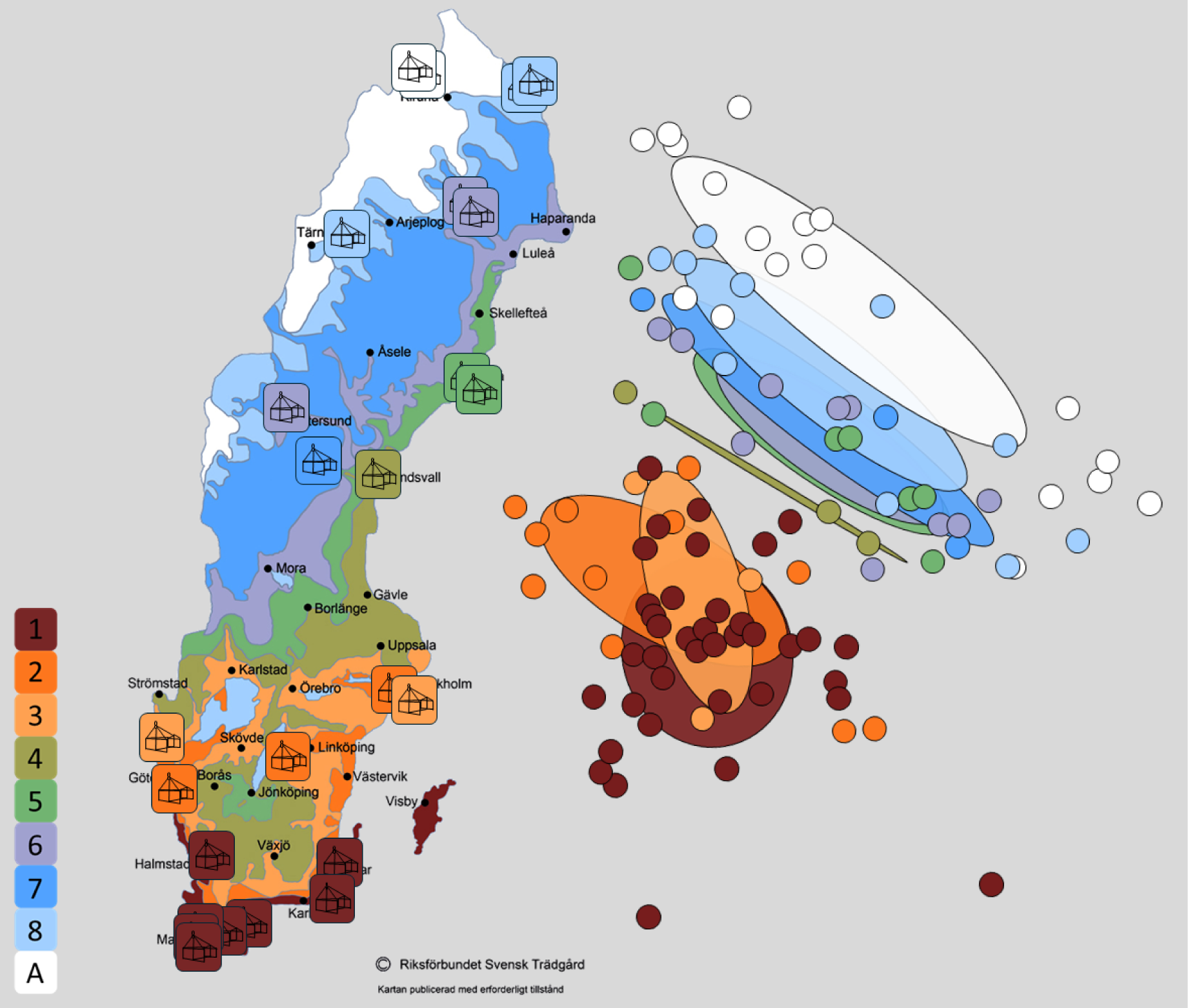
The location of 37 study sites colour-coded according to the plant hardiness zones (odlingszoner, 1-8 and Alpine) of the Swedish Horticultural Society (Riksförbundet Svensk Trädgård) (map used with permission) next to an NMDS plot of study samples colour-coded with the same zones.

**Figure 2.**
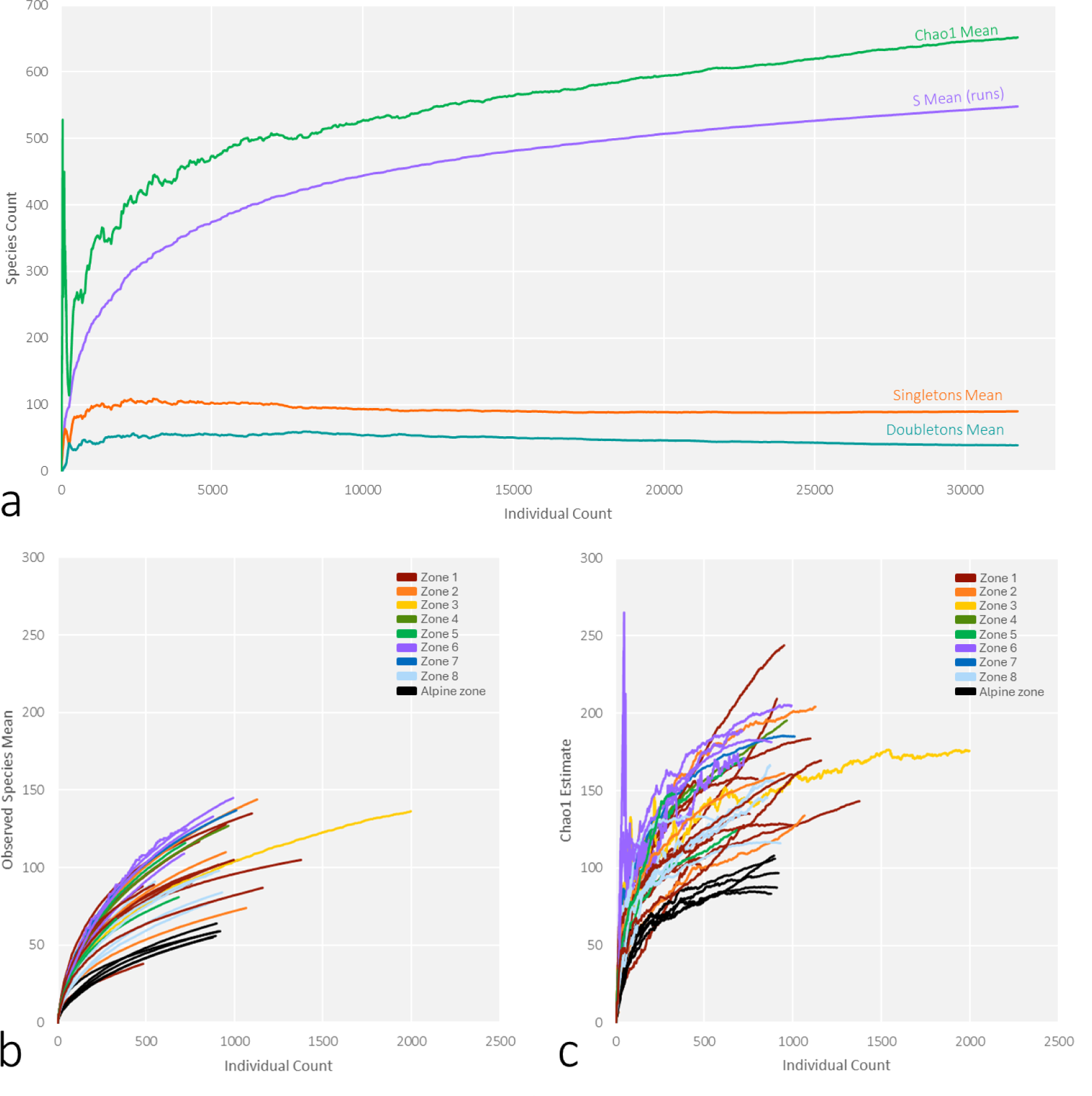
(a) Species accumulation and Chao1 estimate curves for the scuttle fly dataset across Sweden. Notably, current sampling is far from exhaustive, and numbers of singletons and doubletons in our dataset are high regardless of sample size, (b) Species accumulation curves by sampling sites, colour-coded by zone, and (c) Chao1 estimate curves by sampling sites, colour-coded by zone. For a map of the zones using the same colour codes, see Fig. 1.

### Ordinations of community composition

NMDS plots revealed a clear distinction between scuttle fly communities in the southern (zones 1-3) and northern (zones 4-alpine) plant hardiness zones (Fig. 1, Supp. Fig. S5). ANOSIM supported significant differences between southern and northern zones at this threshold (R=0.58, p=0.001), and SIMPER revealed that North-South similarity was just 13.2%, as compared to similarities of 24.6% and 23.2%, respectively, among samples within the northern and southern zones (Supp. Tables S2-S3).

The separation between zones was consistent across clustering thresholds ranging from haplotypes-as-such to a threshold of 1.7% sequence similarity, with stress values around 0.21 (Supp. Fig. S5 top row). Above a threshold of 3%, however, the patterns were increasingly blurred and the stress values were higher (0.25-0.27) (Supp. Fig. S5 bottom row). These patterns were also evident in ANOSIM analyses, where the sample statistic decreased from 0.54 for haplotypes to 0.27 for 5% mOTUs, indicating a decreasing relationship between scuttle fly community composition and plant hardiness zones at higher clustering thresholds (Supp. Table S2). Similarly, in SIMPER analyses, average similarity between zones increased from 20.2% for haplotypes to 41.4% for 5% mOTUs (Supp. Table S3). For species, scuttle fly communities were found to be distinct across most zones (R=0.51, p=0.001), with higher (average=28.5%) similarity within zones than between (average=19.2%) zones (Supp. Tables S2, 3).

Samples from the late spring, midsummer, and late-summer time-periods showed a clear progression along the ranked plant hardiness zones, while offseason samples appeared more randomly distributed (Supp. Fig. S6). Northern sites (IV-alpine) showed higher distinctness across seasons (R=0.59) than when considering all sites (R=0.27) (Supp. Table S4). The between-season similarity of all sites averaged 16.9% and within time-period similarity averaged 23.7% (Supp. Table S5, top). However, for northern sites only (IV-alpine), between time-period similarity was 21.3% and within time-period similarity was 37.2% (Supp. Table S5, bottom).

### Hierarchical Modelling of Species Communities (HMSC)

Models of the four response matrices (species-occurrence, haplotypes-occurrence, species-abundance, haplotypes-abundance) showed relatively good MCMC convergence with potential scale reduction factors of the models’ beta and omega parameters close to one (Supp. Fig. S7).

Explained variance averaged c. 30% for the occurrences and c. 60% for the abundances of both species and haplotypes, but there were large differences in model fit among taxa (Supp. Fig. S8), haplotype occurrence mean ± sd Tjur R^2^=0.30 ± 0.15 (range 0.05 – 0.82); haplotype abundance R^2^=0.61 ± 0.26 (range 0.00-1.00); species presence-absence Tjur R^2^=0.30 ± 0.14 (range 0.05-0.72); species abundance R^2^=0.59 ± 0.26 (range 0.02-1.00)).

Sampling time-period explained the largest fraction of variance, on average, in all four HMSC models (mean 10-11% in the occurrence models; mean 15-16% in the abundance models; Fig. 3, Supp. Fig. S8). Mean annual temperature explained almost as much variance as sampling time-period in the occurrence models (mean 7% and 8% in the species vs haplotype models), but clearly less than sampling season in the abundance models (mean 6% in the species models and 5% in the haplotype models) (Fig. 3, Supp. Fig. S8). All response variables showed strong climatic (including temporal) community structure and there was also strong spatial structure in species and haplotype occurrences linked with the latitudinal temperature gradient across Sweden. Tree cover explained more variance in the abundance models (mean 6% for both species and haplotypes) than in the occurrence models (mean 1% for both species and haplotypes). Trapping effort also explained more variance on average in the abundance models (mean 7% and 8%) than in the occurrence models (mean 1.5% for both), as did the effect of having (vs not having) sequenced the full trap sample (2% mean for occurrence, 6% mean for abundance).

**Figure 3.**
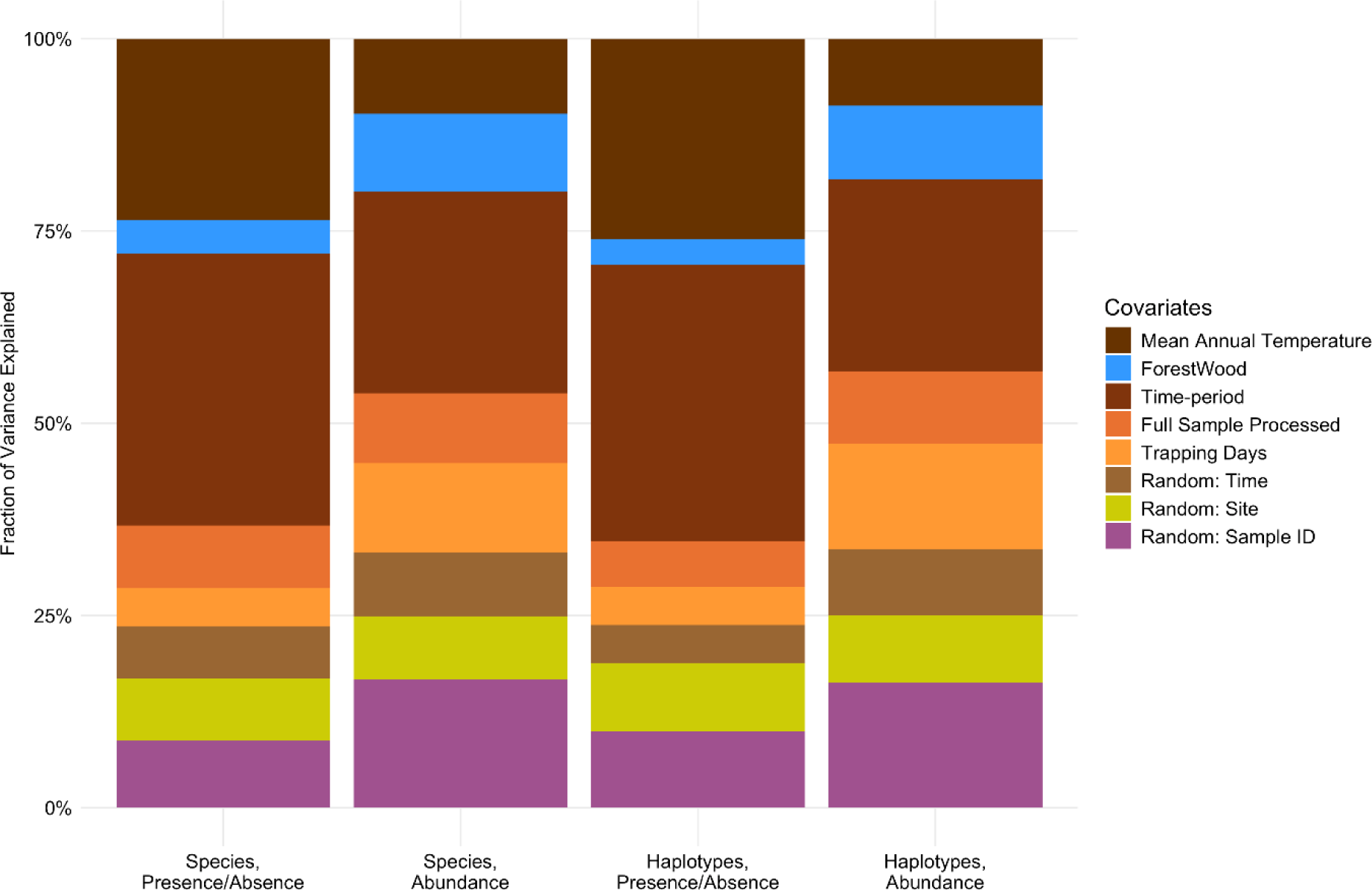
Summary of explained variance in the occurrences and abundances of scuttle fly haplotypes and species across samples (bar height corresponds to meanTjur’s R^2^ across the modelled taxa in the occurrence models and to mean R^2^ in the abundance models). All four models show strong climatic structure (in shades of brown) on scuttle fly communities, as captured by fixed variables describing sampling season and mean annual temperature and a random effect based on median sampling date. The spatial fraction of explained variance (lime green) reflects community structural differences among sites that were not captured by the fixed effect covariates. Differences in sampling effort, i.e. whether or not trapped flies were all sequenced or not and the number of field trapping days per sample, also affect the predictability of community structure (shades of orange). The abundance of species and haplotypes, and to a lesser extent their occurrence, was also strongly structured by habitat type as described by forest and woodland cover (blue). Finally, we included a categorical random effect representing sample identity (purple).

While the occurrences of taxa and haplotypes showed statistically supported responses to all model covariates, their abundances showed responses less frequently, and responses with strong support were primarily with seasonal covariates (Fig. 4).

**Figure 4.**
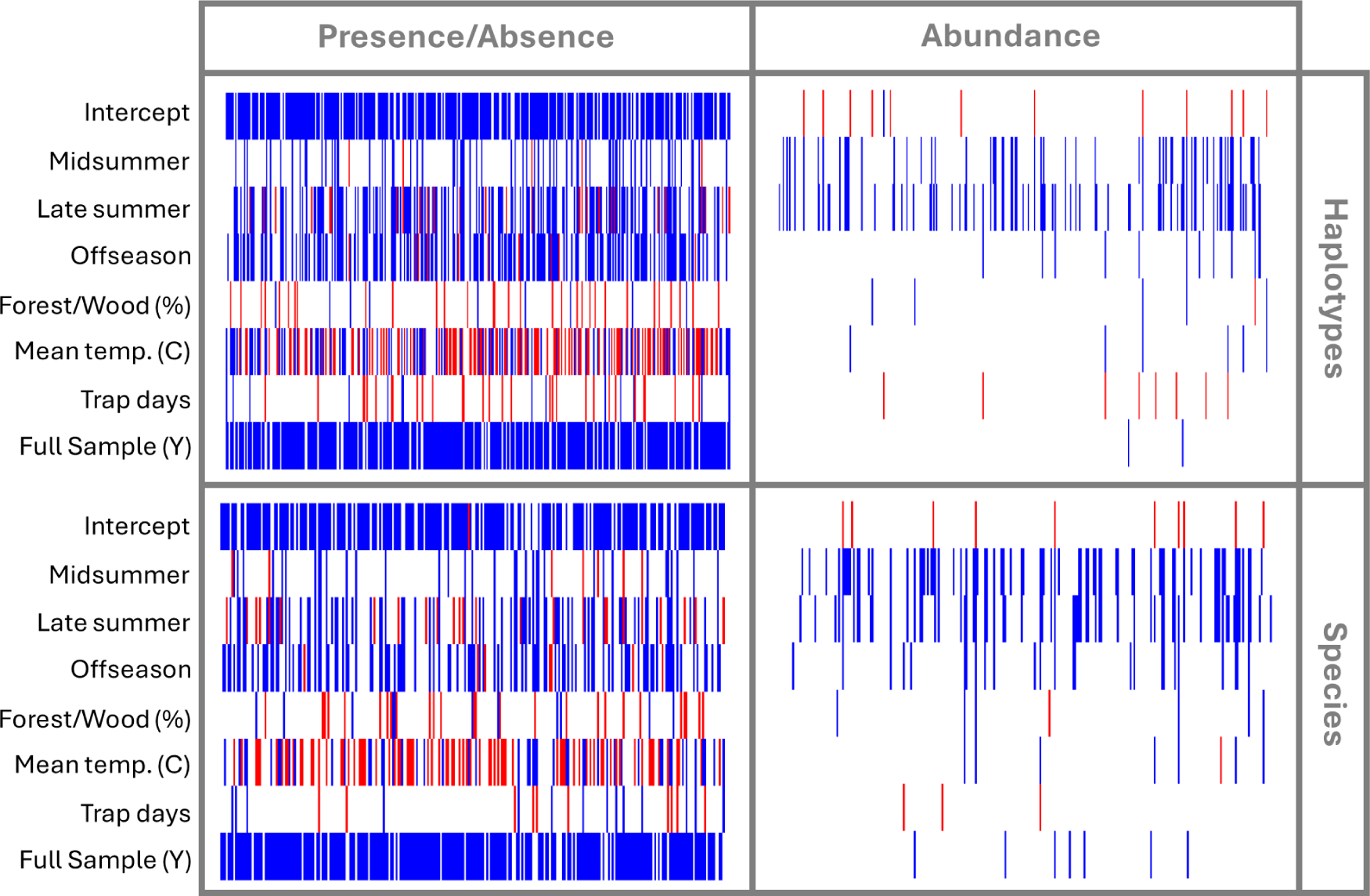
Predicted occurrence and abundance responses of 193 scuttle fly haplotypes and 162 mOTUs (“Species”) to HMSC model covariates. Positive (red) and negative (blue) estimated responses with a posterior probability of 0.95 are illustrated. The three rows below the intercept illustrate the estimated effects of three levels of a categorical variable representing sampling time-period (midsummer, late summer, offseason) relative to the baseline (late spring). The subsequent three rows represent responses to % forest or woodland cover and mean annual temperature (“bio1”) at sampling sites. The final two rows represent the effects of two sampling-related differences among samples: the number of trapping days and whether all specimens in the trap sample were sequenced, and hence available for HMSC analysis, or not.

Most species and haplotypes with a statistically supported seasonal abundance trend peaked in the late spring. Compared to late spring, the midsummer fauna showed a reduction of 28% in species counts and 16% in haplotype counts, respectively, with a further reduction of 24% in both species counts and haplotype counts towards the late summer. Taxa and haplotypes were also usually more prevalent in late spring than in mid- or especially late summer. However, a minority of species and haplotypes (9% and 7%, respectively) showed the reverse pattern, being more prevalent in late summer than in the spring. Offseason captures in the late fall to early winter were consistently low, and the number of offseason samples included in the models was the smallest (n = 20 samples). Nonetheless, 33% of the species modelled and 29% of the haplotypes modelled were occasionally detected in offseason samples.

Taxon occurrences showed a mixture of positive (29% vs 30% for species and haplotypes, respectively) and negative (18% vs 23%) responses to the annual temperature gradient (“bio1”) across Sweden (Fig. 4), reflecting the broad-scale compositional changes from southern towards northern Sweden seen in the NMDS ordinations. Where detected, the occurrence responses of taxa to forest or woodland cover were more often positive than negative (10% positive vs 4% negative for species; 8% vs 2% for haplotypes; Fig. 4). Longer trapping periods did not consistently result in higher detection probabilities of taxa, presumably due to seasonal differences in trapping duration (4% positive vs 5% negative responses for species; 8% positive vs 4% negative for haplotypes; Fig. 4). As expected, most taxa (81-82% in both the species and haplotype models) showed a negative occurrence response to the binary variable indicating whether all specimens in a sample were sequenced or not (“FullSample”) (Fig. 4). The mean annual temperature gradient across Sweden was predicted to affect the prevalence of 38% of species, but does not appear to be a main driver of species abundance (Fig. 4). The predicted effect of the temperature gradient on species prevalence during late summer was positive in 22% of taxa and negative in 16% of taxa (Supp. Fig. S9). Beyond spatial patterns explained by these climatic predictors, there was evidence of localised spatial autocorrelation in species and haplotype site occupancies at scales of less than 40 km. Neither species nor haplotype abundances were spatially autocorrelated, nor did we detect statistical support for temporal autocorrelation in any model.

Residual correlations in the distributions of taxa were detected among sites and samples in both the species and especially the haplotype occurrence models (Supp. Fig. S10). Residual associations of taxa over time were also evident, but less frequently (Supp. Fig. S10). Residual associations between taxon/haplotype occurrences likely indicate that our models either lack or imperfectly represent some of the variables that structure the occurrences of scuttle fly taxa and haplotypes in space and time. No residual associations among taxa were evident among sites or over time in the species or haplotype abundance models, and very few were detected among samples (Supp. Fig. S10). Hence, covariance in the abundances of taxa and haplotypes across occupied samples was well modelled by the environmental and other covariates in these models.

The observed richness of the excluded rare vs modelled common species and haplotypes was strongly positively correlated (R = 0.59 for species and R = 0.80 for haplotypes). Compositional differences between samples in terms of the rare vs common species and haplotypes were also positively correlated (for taxon presence-absence Mantel R = 0.31 for species and Mantel R = 0.64 for haplotypes; for taxon abundance Mantel R = 0.30 for species and Mantel R = 0.60 for haplotypes).

### Community Dissimilarity

Turnover accounted for the bulk of spatial, temporal and spatio-temporal variation in community dissimilarity (Fig. 5). In the spatial analyses, the mean turnover of species and haplotypes between sample pairs was 0.75 and 0.87, respectively, while the corresponding mean values of nestedness were 0.07 and 0.03. In temporal analyses, similarly, the mean values of turnover were 0.45 and 0.74 for species and haplotypes respectively, and the corresponding means of nestedness were 0.13 vs. 0.07. Finally, mean turnover values for the spatio-temporal analyses were 0.75 and 0.84 vs a mean nestedness of 0.07 and 0.04 for species and haplotypes, respectively.

**Figure 5.**
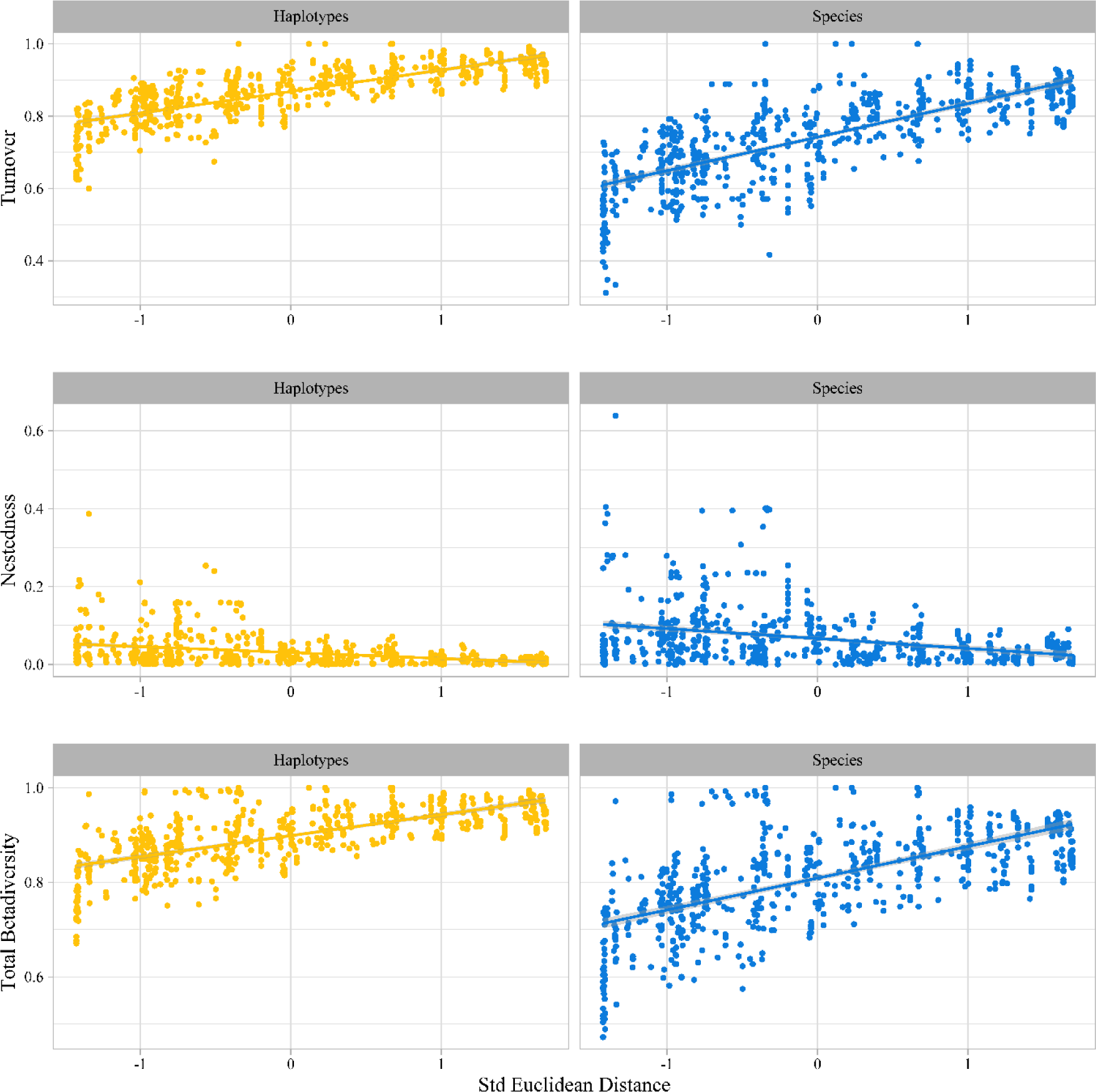
Pairwise community dissimilarity among scuttle fly species and haplotype samples, Following Baselga & Orme (2012), we partitioned overall dissimilarity into its turnover and nestedness components and illustrate these as well as overall dissimilarity as a function of distance (standardized Euclidean distance calculated from sample site coordinates).

Regardless of the time-period and operational taxonomic units used, we found a significant positive correlation between turnover and geographical distances, meaning that communities in closer proximity to each other are more similar in composition (Supp. Fig. S11). The mean spatial turnover of species communities was highest in the summer time periods (midsummer: 0.82, late summer: 0.84) and lower in late spring and offseason (0.78 and 0.72, respectively). However, we did not find any significant correlation between temporal distances (difference in mean week) and turnover (see Table S6). Patterns of nestedness showed no detectable correlation with any of the distances explored (except for the grouped temporal distance) (Supp. Table S6). Within each time-period, scuttle fly communities become more distinct from neighbouring communities from late spring to late summer (Supp. Table S6 and Supp. Fig. S11-13).

## Discussion

Our study marks the first country-wide examination of a dark taxon’s diversity and distribution. It revealed more than 500 species based on processing ca. 31 800 specimens. The ecological analyses suggest that climate change may have profound (and quickly apparent) effects on communities of scuttle flies, that may serve as early indicators of future shifts in the environment. We demonstrate that armed with recent advancements in sequencing technologies, bioinformatics pipelines and molecular barcoding workflows (Meier et al. 2006; Wang et al. 2018, 2018a; Srivathsan et al. 2019b, 2021; Yeo et al. 2020; Hartop et al. 2021), we are now able to resolve patterns in the alpha diversity, spatial and temporal turnover, and species communities of challenging dark taxa. We show that for the scuttle flies in Sweden, a major part of all species remains undiscovered, that local communities show major turnover in space and time, and that ecological patterns are largely robust to the finer details of species delimitation. Below, we will address each of these findings in further detail – and the importance of these patterns in dark taxon biology to biodiversity science.

### Diversity

Our sample only scratched the surface of the scuttle fly fauna of Sweden (Fig. 2, Supp. Figs. S3, S4). While the true Swedish fauna of scuttle flies is still hard to estimate, the 549 putative species found (and 652-713 predicted based on current sampling) greatly exceed the 374 previously documented from the country. Previous estimates for the scuttle fly fauna have ranged from 1 100 to nearly 2 000 species, suggesting that the true numbers may be even higher (Ronquist et al. 2020). This would not be surprising, given that our study included sampling from just 37 traps in a country of over 450 000 square kilometers; i.e., many habitats remained unsampled. However, the available data confirms that for dark taxa the basic pattern in ecology holds that most species are rare and that finding all species would require massive sampling (Callaghan et al. 2023).

Where the remaining species will be lurking is unknown. While Malaise traps are remarkably effective at capturing flies (Karlsson et al. 2020; Skvarla et al. 2020), other trapping methods would certainly reveal additional species. An “All Diptera Biodiversity Inventory” conducted in Costa Rica utilised a wide variety of methods (Borkent and Brown 2015), revealing that 59% of fly species were unique to a specific collection method (Borkent et al. 2018). Intriguingly, species-rich taxa with particularly interesting and often impactful biologies (e.g. parasitoids) may be underrepresented in Malaise trap samples. For example, Brown (1996) reported that nearly half of the species of ant-parasitising scuttle flies were caught exclusively over army ant raids. Similar results were obtained by Disney et al. (1982), who found some species of scuttle flies were uniquely or primarily collected in water traps rather than Malaise traps. Another possible frontier is the canopy. Ongoing research in upper levels of the canopy in the Amazon suggests that it contains many scuttle fly species not found in lower sampling elevations (Gilliland 2021), contrasting with previous data showing scuttle flies as a taxon primarily collected at ground-level (Kitching et al. 2005). In a limited sample (159 specimens) from canopy trap prototypes run concurrently with the Malaise trap sampling for this study we found a single unique mOTU not seen in the 31 794 scuttle flies from ground level, indicating that even in a temperate environment we may have more to find in the canopy.

Thus, no single sampling method will suffice to reveal all taxa and additional trapping sites that target regions or habitats un- or under-represented in the current study will also be needed for an exhaustive inventory of scuttle flies. Some of this complementary sampling may have already been carried out by the Swedish Malaise Trap Project years ago (Karlsson et al. 2020). An excellent follow-up to this study would be sequencing scuttle fly samples from that sampling campaign, or from another insect campaign that is ongoing in Sweden – the Insect Biome Atlas project (www.insectbiomeatlas.com).

In addition to focusing future efforts on revealing more species, the analysis of other life stages may offer a more nuanced understanding of scuttle fly communities. Our study is based on a sample of adult flies. Patterns for the longer-living larval stages are at this point unknown, as they are not easily collected.

### Lessons from Ecological Analyses

Our ordinations of community composition suggested strong structuring of scuttle fly communities by Swedish horticultural zones (Fig. 1) and season (Supp. Fig. S6). Clear north-south structuring of the Swedish insect fauna has also been observed in damselflies, mosquitos, and caddisflies (Gullefors 2008, Wellenreuther et al. 2012, Lundström et al. 2013), and strong phenological patterning is a hallmark of any high-latitude fauna (Wolda, 1988). Our ordinations also indicated that spatial structuring was largely independent of mOTU clustering threshold at or below the species proxy level (Supp. Fig. S5).

To address these findings more rigorously and without the constraints of pre-determined zones, we implemented HMSC. This confirmed that scuttle fly community variation is both highly seasonal and strongly tied to the latitudinal temperature gradient over Sweden, with all models strongly predicted by climatic covariates (Fig. 3, Supp. Fig. 8). The presence of significant spatial autocorrelation in the HMSC occurrence models reflects the gradation of scuttle fly distributions across space. Scuttle fly communities also showed clear compositional changes over time during the warm season, from late spring through mid and late summer, while offseason sampling was too inconsistent to reveal any patterns (Supp. Figs. S2, S6). Consistent with our findings that scuttle fly communities are driven by climatic covariates, we found more rapid spatial turnover in taxa at the species than at the haplotype level (Fig. 5). Recent work has predicted that when dispersal limitation is the dominant driver of species distributions, the rate of spatial turnover of biological communities will be similar at both the haplotype and species levels (Baselga et al. 2022). Conversely, when environmental conditions strongly constrain species ranges, community similarity is predicted to decay at different rates across genealogical scales.

Our results have several implications in the face of climate change. Individual taxa show a mixture of positive and negative responses to mean annual temperature, and to a lesser extent to seasonality. This implies that we may see a substantial number of both climate change “winners and losers” in the future, as species ranges and phenology expand, contract, or shift (Fig. 4, Supp. Fig. S9). Adult scuttle flies are ephemeral – with short lifespans and high turnover and mobility – they may respond rapidly and serve as indicators for future shifts of other taxa and in the environment more broadly. Our observation of steeply declining abundance from late spring through the summer into the offseason may partially reflect the extreme temperatures and drought in Europe in summer 2018 (Bakke et al. 2020, Bastos et al. 2020). While this suggests that our seasonal results may be atypical, they are perhaps also a sign of summers to come, as climate change increases the frequency and severity of these events (Hari et al. 2020).

With our initial results confirmed by HMSC, the predictive value of the simple plant hardiness map for scuttle fly distributions offers excellent news. It suggests that even such a relatively coarse-grained tool can be used as a reliable indicator of the regional compositions of scuttle flies. That this is the case is only intuitive: Many scuttle flies exist close to ground level and therefore, like plants, their distributions may be closely tied to soil temperature, acidity, moisture, or composition (all of which would be interesting covariates to explore in future modelling). Additionally, some species are known to have direct interactions with plants, fungi, and ground-dwelling insects – factors which may again tie them to the microclimate at ground level. Future studies might focus on these microenvironmental variables, to address the unexplained variance in scuttle fly communities from this study.

### Dark Taxon Zoology

Dark taxa have historically been largely ignored due to the complexities involved in studying groups of highly diverse and abundant organisms of small size. How, then, do we make the study of Dark Taxa efficient and start “dark taxon zoology” to respond to the need for quantitative data on biodiversity?

To address this, we assessed whether the taxa analysed need to exactly match taxonomically validated species for ecological hypotheses to be addressed. If not, we may avoid endless discussions about species delimitation and can proceed with ecological analyses. Promisingly, our current results suggest that ecological patterns are largely robust to the exact clustering thresholds used. Overall biodiversity patterns, patterns of community turnover and drivers of distribution were virtually identical using thresholds from 0% (haplotype data) to 1.7% (Figs. 3-5, Supp. Figs. S5, 8, 11-13).

Haplotype data are convenient because they do not require any taxonomic decisions, which may please molecular ecologists, with the important caveat that exceeding an appropriate species proxy threshold will obscure patterns because too many species will be lumped together (Supp. Fig. S5, bottom row).

A second potential stumbling block in the study of dark taxa lies in the large numbers of rare species. If understanding basic ecological patterns is dependent on these rare species, dark taxon zoology will be difficult. Our results indicate that both richness and compositional differences between samples of rare versus common species and haplotypes were both positively correlated. This suggests that more common and rarer species respond similarly to the same drivers. Again, this is excellent news, since it suggests that ecological inferences regarding the drivers of species distribution and community composition can be based on the more common species – which are much easier to detect.

## Conclusions

We here argue for a dark taxon zoology and illustrate that it is not a hopeless undertaking by targeting scuttle flies as a typical dark taxon in Sweden. We sample across the entire country, we estimate the species richness, resolve patterns of distribution across time and space, and pinpoint environmental features that drive these distributions. Our results suggest that such assessments will be insensitive to specific taxonomic cut-offs, and robust to undersampling of rare taxa. Overall, the study hopes to contribute to a more quantitative approach to biodiversity. In the future, advances in molecular workflows, bioinformatics, robotics and automation will make these groups increasingly efficient to study (Meier et al. 2024). We hope that our case study will serve to propel forward Dark Taxon Zoology, bringing the main part of diversity into the realm of biodiversity science.

## Materials and Methods

### Target taxon: the scuttle flies of Sweden

Sweden has one of the best-known animal faunas in the world due to efforts dating from Carl Linnaeus to the Swedish Taxonomy Initiative and Malaise Trap Project (Linnaeus 1735; Miller 2005; Karlsson et al. 2020, but see Ronquist et al. 2020). While 374 species of scuttle flies have been documented in the country (SLU Artdatabanken 2021), this is an underestimate. For comparison, a single suburban garden in Cambridge, UK, yielded nearly 100 species of scuttle flies (Brown and Hartop 2017), while backyards in Los Angeles, California, can support up to 82 species (Brown and Hartop 2017). Previous estimates of scuttle fly diversity in Sweden have proposed that the true fauna may approach 2000 species (CNE estimate in Ronquist et al. 2020), but this may be an overestimate based on an error-prone process of morphological identification (Ronquist et al. 2020).

### Sampling

To start resolving the species richness, spatiotemporal distribution, and community composition of Swedish scuttle flies, we sampled communities of flying insects at 37 locations across Sweden (Fig. 1, Supp. Fig. S1, Supp. Table S1). These samples were collected in ∼80% ethanol using Townes-style Malaise traps (Townes 1972). Scuttle flies were sorted from the trap samples and preserved in ∼80% ethanol at −20°C.

Sampling started in May 2018 and continued into the following year. Although trapping was continuous, the sample periods varied across sites due to their extensive distribution across Sweden. Four time periods that matched the summer phenology were therefore established in sequence for each site: late spring, midsummer, late summer, and offseason (Supp. Fig. S2, Supp. Table S1). Some offseason samples could not be retrieved in late 2018 and were collected in 2019. To ensure uniformity in the number of weeks per sampling period and avoid introducing biases, we attributed the latest sampling date for the offseason traps in 2018 to those samples collected in 2019.

Additionally, late spring samples were only obtained from 25 sites, as 12 sites were not installed until later in the season (see Supp. Table S1 for sampling details). To compare richness estimates across seasons, we plotted accumulation curves for each time-period excluding the 12 traps that were not yet installed in late spring.

### Sequencing and bioinformatics

A total of 136 samples were selected for analysis. Most samples (94) contained thousands of scuttle fly specimens and subsampling was needed. Individuals were randomly selected with a minimum of two 96-well microplates of scuttle flies processed per sample. This resulted in a total of at least 190 specimens extracted per site and time-period for most samples. 31 samples containing fewer than 190 specimens and were thus processed completely, and two samples were found to contain no scuttle flies. Additional plates of specimens from nine samples that had been processed in an earlier study (Hartop et al. 2022) were also included (Supp. Table S1). All sample information, including numbers of barcodes obtained per sample and whether a sample was fully processed, is found in Supp. Table S1.

DNA were obtained from the specimens using 10 μL of HotSHOT lysis buffer (Truett et al. 2000). The extraction was carried out in a thermocycler at 65 °C for 18 minutes, then for 2 minutes at 98 °C. Amplification was carried out on a 313-bp fragment of Cytochrome Oxidase 1 (*COI*) using primers m1COlintF: 5’-GGWACWGGWTGAACWGTWTAYCCYCC-3’ (Leray et al. 2013) and modified jgHCO2198: 5’-TANACYTCNGGRTGNCCRAARAAYCA-3’ (Geller et al. 2013). For a small subset of samples, *COI* was amplified using the primer pair jgHCO2198 and LCO1490 (Folmer et al. 1994; Geller et al. 2013).

Amplifications were conducted with tagged primers following Wang et al. (2018) for Illumina and Srivathsan et al. (2019b, 2021) for MinION. PCR reactions contained 4 μl Mastermix from CWBio, 1 μl of 1 mg/ml BSA, 1 μl of 10 μM of each primer, and 1 μl of DNA. PCR conditions were a 5 min initial denaturation at 94 °C followed by 35 cycles of denaturation/annealing/extension (94 °C (1 min)/47 °C (2 min)/72 °C (1 min)), and a final extension at 72 °C (5 min). A subset of wells (N=8-12 with negative control) from each PCR plate was run on an agarose gel to check for plate-wide failure, before products were pooled and then purified using AMPure XP beads (Beckman Coulter Life Sciences, Indiana, USA). DNA concentration was quantified with a Qubit™ dsDNA HS Assay Kit (Invitrogen, California, USA).

Illumina and MinION sequencing were used to sequence the amplicons for this study. Illumina libraries were prepared using a TruSeq DNA PCR-free kit to obtain 250bp PE sequences using Illumina HiSeq 2500; the sequencing was outsourced. Nanopore sequencing using MinION was conducted in house following Srivathsan et al. (2019b). Library preparation was conducted using either SQK-LSK109 or SQK-LSK110 ligation sequencing kits (Oxford Nanopore Technologies, Oxford, UK) with 200-ng pooled and purified PCR products. The manufacturer’s instructions were followed except for use of 1× Ampure beads instead of 0.4× as suggested in the instructions because the minibarcode amplicons in our experiments were short (∼391 bp with primers and tags), and a modified protocol for end-repair as described in Srivathsan et al. (2021). The sequencing was carried out using a MinION sequencer with either R9.4.1 or R10.3 flow cells for a maximum of 72 hours. Basecalling was conducted using Guppy versions 2.3.5+53a111f and 4.2.3+f90bd04. FastQ files were demultiplexed to obtain individual *COI* sequences identifiable back to individual specimens with unique specimen codes. This was done following Srivathsan et al. (2019a, 2021) for MinION and Wang et al. (2018) for Illumina.

To exclude contaminants and potential mis-sorts, *COI* sequences were matched using BLAST to NCBI GenBank’s nucleotide database and sequences with >97% similarity to non-scuttle fly taxa were removed. Sequences were then aligned using MAFFT v7 (Katoh and Standley 2013) and clustered into molecular Operational Taxonomic Units (mOTUs) using Objective Clustering (part of “TaxonDNA”, see Meier et al. 2006), a distance-based method that groups sequences based on a user-defined threshold for minimum interspecific uncorrected p-distance.

A species-proxy clustering threshold of 1.7% was used for the primary analyses as this was the distance that maximized cluster congruence to species-level morphology for a subset of 18 000 scuttle flies from this dataset (Hartop et al. 2022). Hereafter, references to “species” are to mOTUs at this threshold. To assess the impact of the selected threshold on distribution patterns, we examined other thresholds ranging from 0-5% and ran models using both species and haplotype data.

### Diversity

To characterise local and national species richness, we used accumulation curves and Chao1 richness, as implemented in EstimateS (Colwell 2013) and R (R Development Core Team) package iNEXT (Chao et al. 2014) for total diversity, diversity per zone, and diversity per site. Following the approach of Ronquist et al. (2020), a combined non-parametric estimator (CNE) was used to obtain an alternative estimate of total species richness.

### Characterisation of community variation

To visualise scuttle fly community composition in the context of geographic and climatic variation, we used Non-metric Multi-Dimensional Scaling (NMDS) plotted with R packages ggplot2 (Wickham, 2016) and Vegan v2.5-4 (Oksanen et al., 2019) using both abundance-based (Bray-Curtis) and incidence-based (Jaccard) dissimilarity indices. Since both metrics yielded nearly identical results, the results in the paper are based on Bray-Curtis indices. Sites were classified according to the nine plant hardiness zones of the Swedish Horticultural Society that synthesize climatic variables of particular importance to horticultural plants into horticultural zones: strong and rapid temperature changes, very low temperatures sustained for long periods, evaporation occurring when the sun is shining but the ground is still frozen, and temporal considerations of climatic and environmental conditions (Fig. 1, Riksförbundet Svensk Trädgård 2018, used with permission). As these zones have proven useful in identifying the survival and growth of horticultural plants across Sweden, we hypothesized that they may offer a relevant description of the environment for any organism sensitive to similar climatic variables – including scuttle flies. They also offer a more regional approach than the broad biogeographic classification previously used for spatial analysis of scuttle flies in Sweden (Ronquist et al. 2020). We therefore tested whether the zones could predict scuttle fly richness and distributions (Disney 1994; Kitching et al. 2005; McGlynn et al. 2019).

To quantify differences in scuttle fly community composition between plant hardiness zones and to assess the significance of those differences, we used Analysis of Similarities (ANOSIM) and Similarity Percentage analysis (SIMPER). These tests were run with PRIMER v7 (Clarke and Gorley 2006).

We excluded samples that contained fewer than 100 specimens from the ordination analyses, as small sample sizes can artificially generate large distances in the NMDS plots, thus obscuring structured variation in community composition arising from responses to the environment.

### Hierarchical Modelling of Species Communities

To relate variation in community composition to continuous environmental drivers without the assumptions of zones used in our NMDS visualisations, we used Hierarchical Modelling of Species Communities (HMSC; (Ovaskainen et al. 2017; Ovaskainen and Abrego 2020), a type of joint species distribution model (Warton et al. 2015)). Due to the zero-inflated nature of the data, we fitted hurdle models, i.e., one model for the occurrence of taxa across samples (probit regression), and a second model for their abundance conditional on presence (linear regression for log-transformed count data, with zeros masked as missing data), henceforth referred to simply as an “abundance model”.

To evaluate the impact of the criterion used in species delimitation, we used four different response variables: species presence-absence, species abundance (where present), haplotype occurrence and haplotype abundance (where present), thus resulting in four HMSC models being fit. As these analyses are uninformative for taxa with very sparse data, we included only haplotypes or species with at least five occurrences (n = 391 haplotypes and 273 species, out of the 2 697 haplotypes and 549 species observed; see Results). To be able to relate each sample to specific climatic conditions, we also excluded 15 samples for which the trapping duration exceeded approximately a month (>34 days).

We included five predictor variables, coded as fixed effects, in our HMSC models. To test for seasonal changes in scuttle fly communities among the sampling time-periods, we included the four-category variable “Time.Period”. To assess the effect of the spatial gradient in mean climatic conditions across Sweden on scuttle fly communities, we included the continuous variable mean annual temperature (“bio1” from the Worldclim database, Fick and Hijmans 2017). The importance of tree cover for scuttle fly distributions was assessed by including a continuous variable describing the percentage of forest or woodland cover within a 50m buffer (“ForestWood”) derived from the Swedish National Land Cover Database (https://www.naturvardsverket.se/en/services-and-permits/maps-and-map-services/national-land-cover-database/). Site values for “bio1” and “ForestWood” were extracted using the R package raster (Hijmans 2024). To account for the effect of sampling effort on taxon detection, we further included a continuous variable quantifying the total number of trapping days per sample (“TrapDays”) and a binary variable indicating whether all specimens in a sample were sequenced or not (“FullSample”). The latter, binary variable defines whether the sample was fully sequenced or not, and was included to account for the higher likelihood of encountering taxa in samples that had very high individual abundances (i.e., samples that could not be fully sequenced).

We moreover included a spatially explicit random effect based on sample site coordinates (“Random: site”) and a temporally explicit random effect based on median sampling date (“Random: time”) to model any spatial or temporal autocorrelation in the scuttle fly dataset, and a categorical random effect representing sample identity (“Random: sample”).

We fitted the model with the R-package HMSC (Tikhonov et al. 2020) assuming the default prior distributions. We sampled the posterior distribution of four MCMC chains, each of which was run for 37 500 iterations, of which the first 12 500 were removed as burn-in. The iterations were thinned by 100 to yield 250 posterior samples per chain, and thus 1 000 posterior samples in total. To explore the rate of Markov chain Monte Carlo (MCMC) convergence, we also fitted otherwise identical models but with 375 iterations (burn-in 125, thin 1) and 3,750 iterations (burn-in 1 250, thin 10).

Convergence was assessed by examining the potential scale reduction factor (PSRF) distribution over the fixed effect (*β*) and random effect (*Ω*) parameters, equivalent to the Gelman–Rubin statistic (Gelman et al. 2014).

We examined the explanatory power of the models through species-specific coefficients of discrimination (Tjur R^2^) for the occurrence models, which measure how well the model discriminates those samples in which a taxon occurs from those in which it does not occur, and R^2^ for the abundance models. Tjur R^2^ is defined as the difference in average model-predicted probability of occurrence for samples in which the species is present vs absent (Tjur 2009).

To assess whether the distributions of rare taxa are likely to be driven by similar factors to the common ones, we calculated Pearson and Mantel correlations, respectively, between the observed species richness and community composition (Bray-Curtis dissimilarity) of the modelled more common taxa and the excluded rare taxa (those with <5 occurrences).

### Community dissimilarity

To further partition variation in scuttle fly community composition in time (between seasons) and space (across Sweden), we dissected overall community dissimilarity (β-diversity) into its turnover (i.e. species replacement) and nestedness (differences of species richness between sites) components (Baselga, 2010).

Our analyses encompassed three aspects: 1) spatial differences; differences in community composition between each sampling site considering their geographical distance, 2) temporal differences; differences in community composition between each sampling period, accounting for their temporal distance in weeks, and 3) spatio-temporal differences; community differences between each sampling site and sampling period including their joint effect. We computed pairwise Jaccard community dissimilarity values using both the species and haplotypes datasets for comparison. Given the observed geographical distance between sampling sites, these distances were rescaled to a mean of zero and a standard deviation of one before analyses. To characterise the temporal difference between samples in each sample pair, we used the difference in the mean week of sampling (for sample-specific details, see Supp. Table S1). Values of total β-diversity, turnover, and nestedness were calculated for each pairwise comparisons of sampling sites, sampling period and sample pairs. We excluded self-pairs and included data for each pair only once. To test the correlation between each beta diversity component and the distances in space and time we used Mantel tests based on Pearson moment correlations. All calculations were implemented in R package ‘betapart’ (Baselga & Orme, 2012).

## Declarations

### Availability of Data and Materials

Data and scripts are available on the project GitHub page at https://github.com/leshonlee/DocumentingPhorids. General HMSC pipeline scripts are available at https://www2.helsinki.fi/en/researchgroups/statistical-ecology/hmsc.

### Competing Interests

The authors declare that they have no competing interests.

### Funding

EH was funded by Swedish Taxonomy Initiative grant 2016-203 4.3. TR and OO were funded by the European Research Council (ERC) under the European Union’s Horizon 2020 research and innovation programme (ERC-synergy grant 856506—LIFEPLAN). OO was funded by Academy of Finland (grant no. 336212 and 345110). MJ was supported by the Academy of Finland’s ‘Thriving Nature’ research profiling action.

### Authors’ Contributions

EH and RM designed the study. EH and LL generated the data. EH, LL, AM, MJ, and PPA analysed the data. EH, LL, and RM wrote the initial draft of the manuscript. All authors contributed to manuscript revision, read, and approved the submitted version.

## Acknowledgements

We thank the Scuttle Fly Sorting Party crew at Station Linné – Carina Romero Ugarph, Harald Havnås, Johan Ennerfelt, Marianne “Mia” Blomqvist, Nino Pettersson, Robert Ennerfelt, and especially Dave Karlsson. We thank the members of the Evolutionary Biology Lab at the National University of Singapore for help with the many hours of wetlab work. We thank Darren Yeo for assisting us with visualisations. We thank Inger Ekrem at Riksförbundet Svensk Trädgård for helping with the plant hardiness zone map, Tomas Lagerström for further information on the map, and Eva Ronquist for first bringing our attention to the map. We thank the Swedish Taxonomy Initiative for the support to investigate the scuttle flies of Sweden, this study brings us one step closer to the ultimate goal of describing all of these species.

## Notes

https://github.com/leshonlee/DocumentingPhorids.

### Competing Interest Statement

The authors have declared no competing interest.

### Summary of Updates

Author affiliations updated, minor revisions have been made to format and figure numbering.

