## Supplemental Materials for "RESOLVING BIOLOGY’S DARK MATTER: SPECIES RICHNESS, SPATIOTEMPORAL DISTRIBUTION, AND COMMUNITY COMPOSITION OF A DARK TAXON"

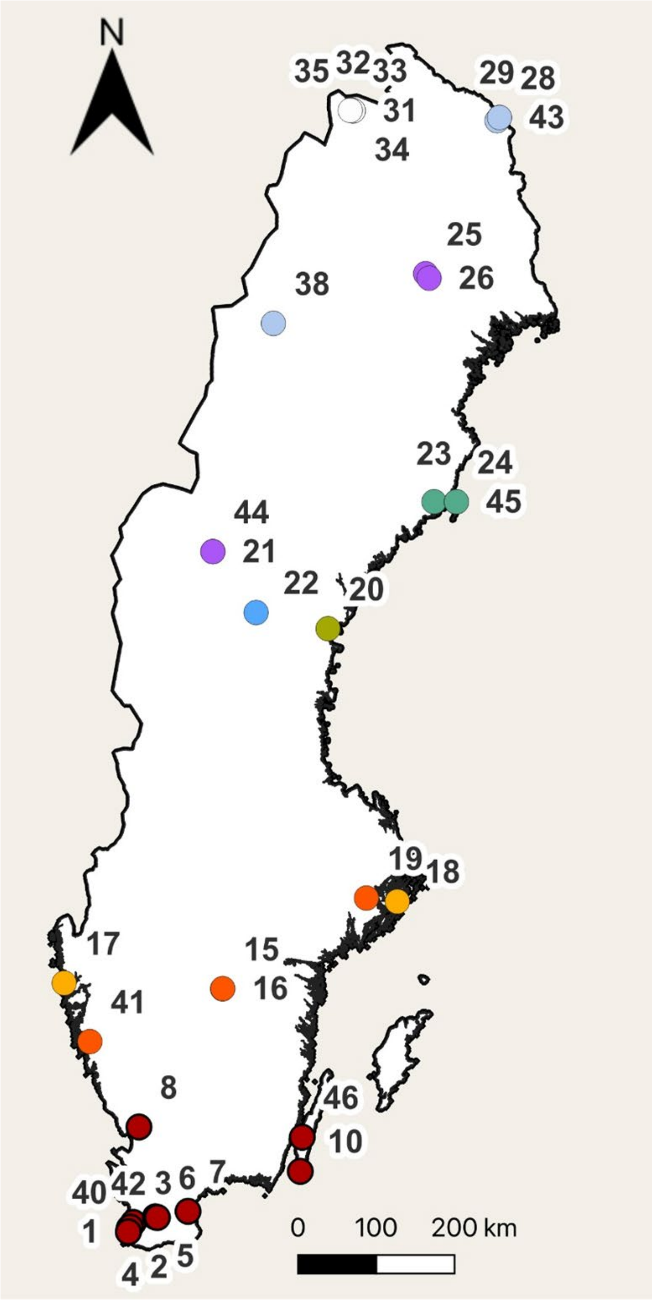


Supplementary Figure S1. Map of study sites.

*A map of the 37 study sites at which scuttle flies were sampled, colour coded according to Swedish horticultural zones (see Fig. 1).*

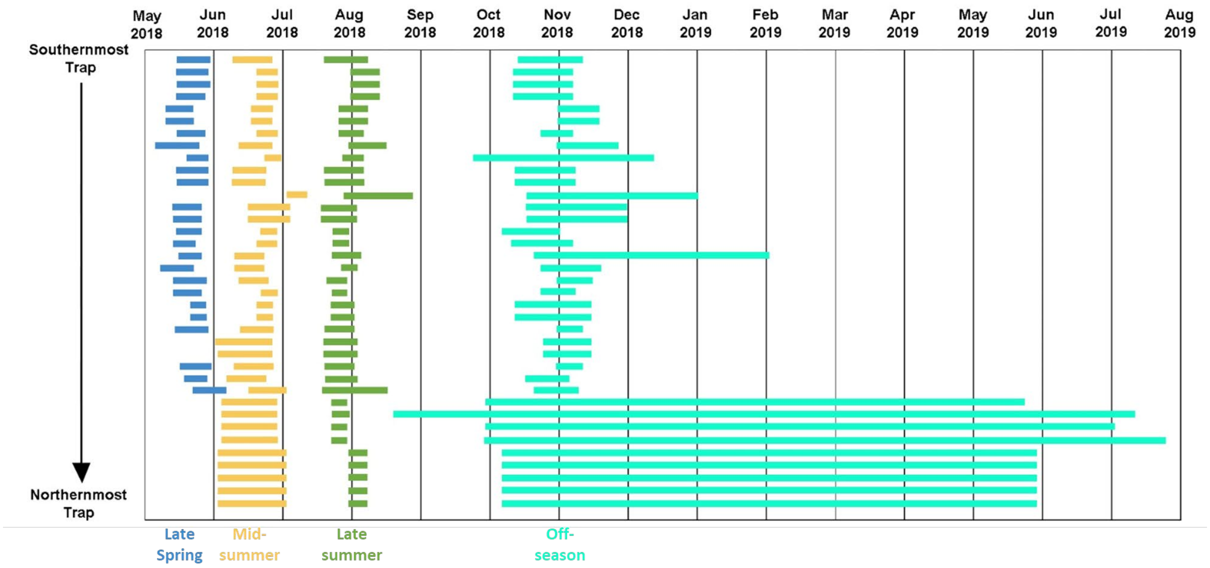


Supplementary Figure S2. Timeline of analysed samples.

*Four sequential samples from each site were selected to represent scuttle fly communities in the late spring (mid-late May, in blue), midsummer (June, in yellow), late summer (late July – early August, in green) and the offseason (last sample of the year, in teal). Many of the traps in the north are missing late spring samples as they were not set up until later in the season. These same sites have long offseason samples as they were not collected until the following spring.*

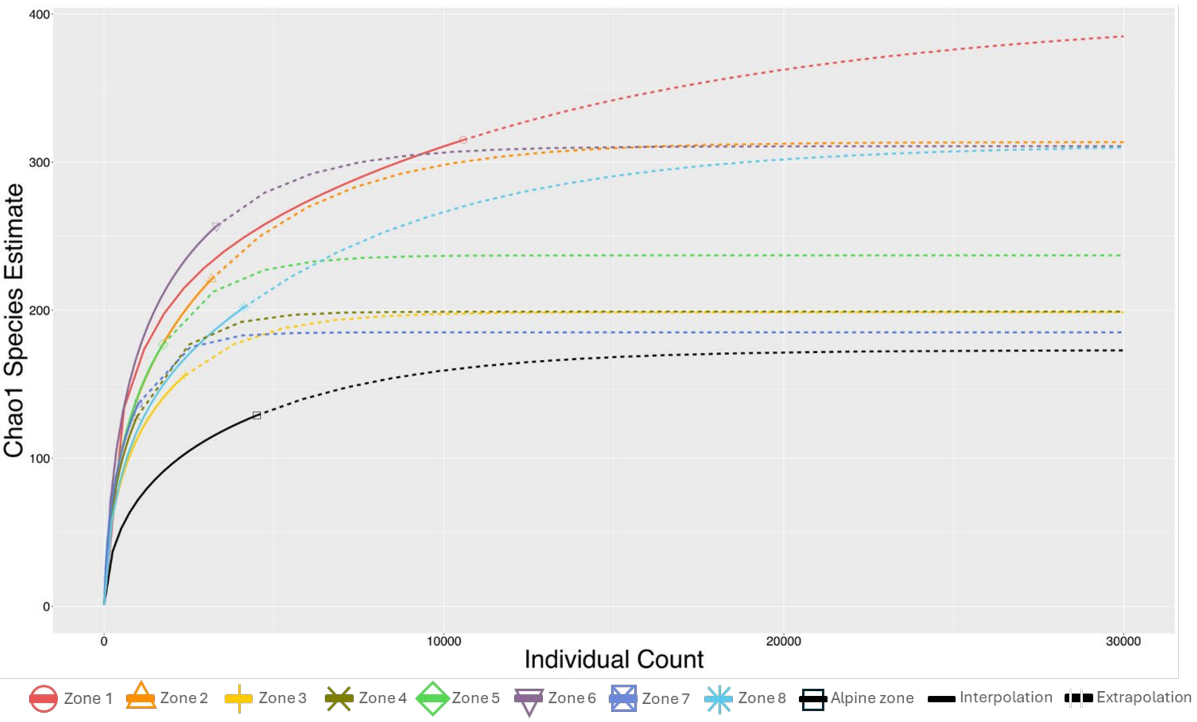


Supplementary Figure S3. Chao1 estimates of scuttle fly species richness per zone.

*Colour-coding is according to plant hardiness zones; for a map of the zones using the same colour codes, see Fig. 1.*

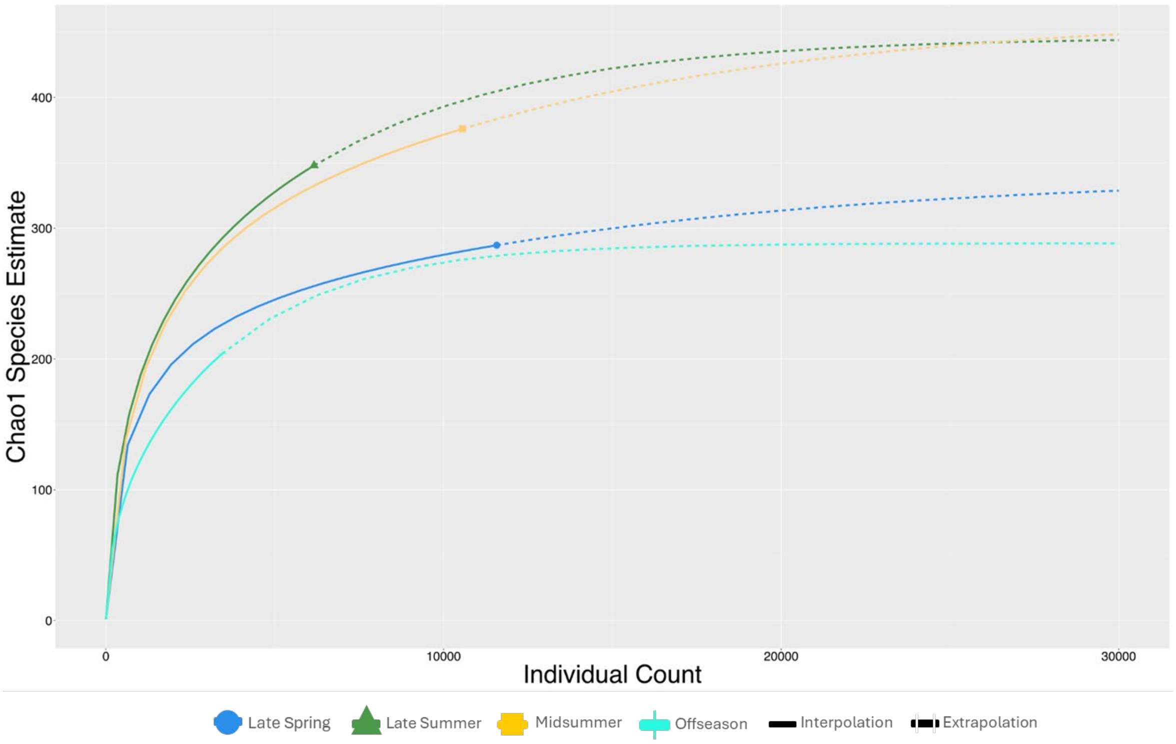


Supplementary Figure S4. Species accumulation curves of scuttle flies by season.

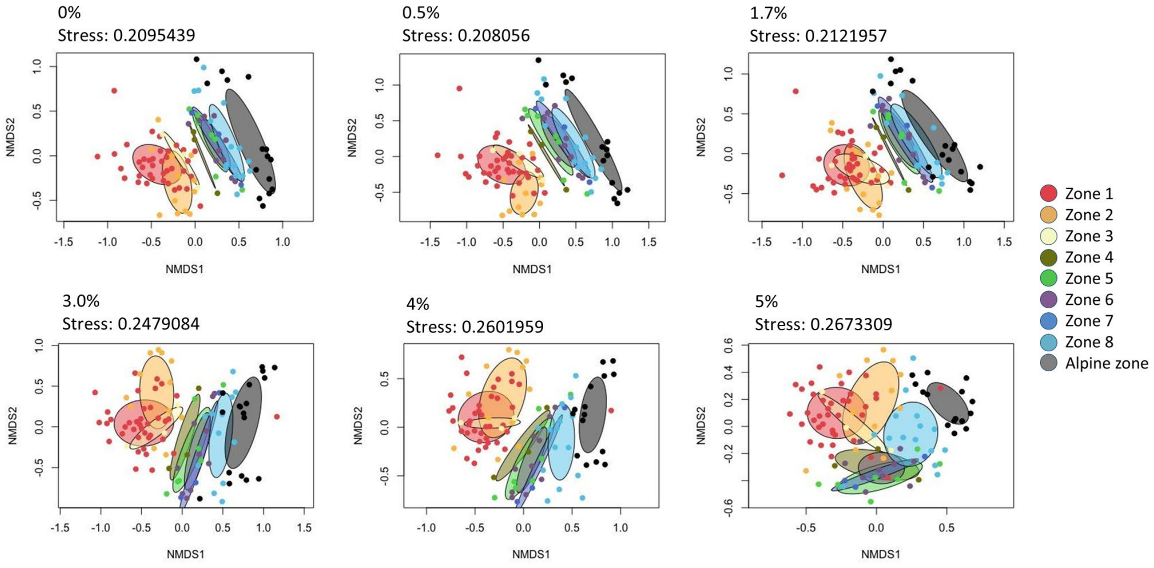


Supplementary Figure S5. Consistency in community patterning, as based on different clustering threshold for species delimitation.

*Shown are NMDS plots with scuttle fly mOTUs calculated at different thresholds with ellipses coloured according to plant hardiness zones. The pattern is consistent from 0-1.7%, after which there is a clear shift and as the threshold increase, the pattern begins to blur as species are lumped. Colour-coding reflects plant hardiness zones, shown in the same colours in Fig. 1.*

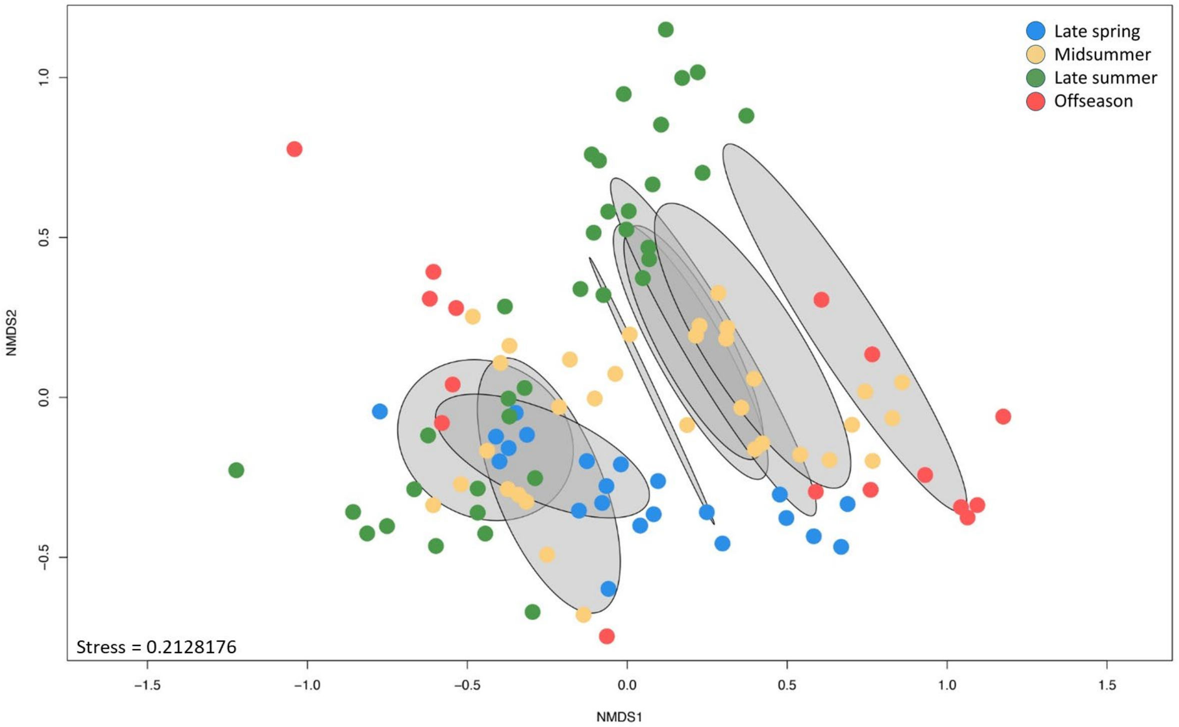


Supplementary Figure S6. NMDS plot of all samples (species, threshold of 100 specimens) with samples colour-coded according to time-period.
*Except for the "Offseason" period, seasonal catches run linearly through zonal ellipses, in grey.*

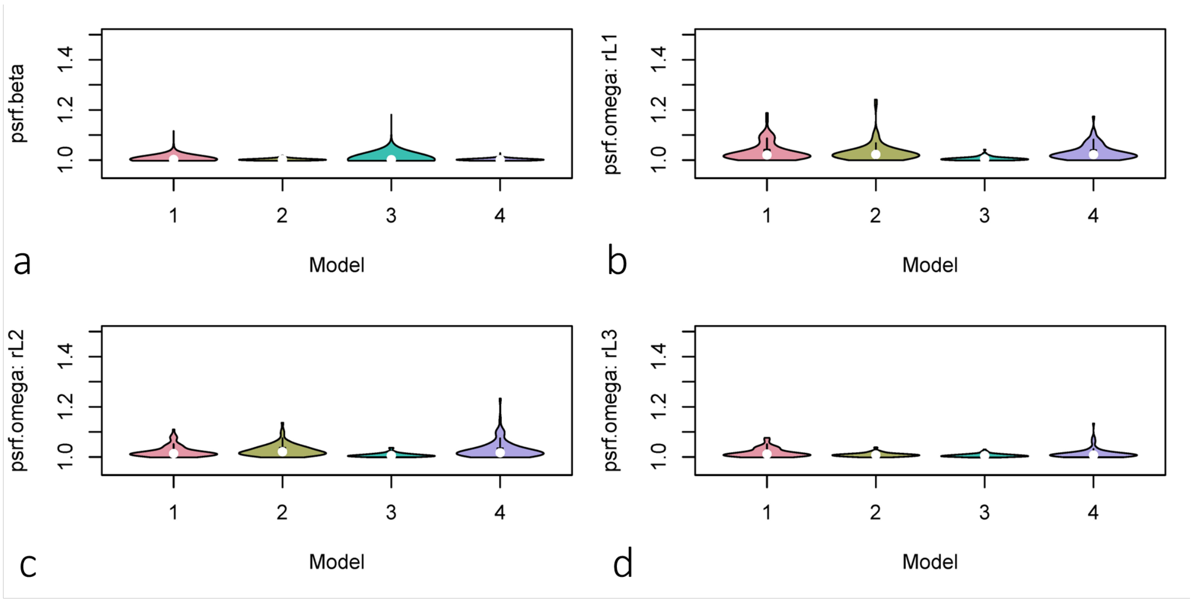


Supplementary Figure S7. Violin plots showing potential scale reduction factors (PSRF) for the beta and omega parameters of four HMSC models: species occurrence (1), species abundance (2), haplotype occurrence (3), and haplotype abundance (4).
*Shown are the PSRF values for (a) fixed effect (β) parameters, (b) random effect (Ω) parameter space, (c) random effect (Ω)* ***parameter*** *year and (d) random effect (Ω) parameter sample-level.*

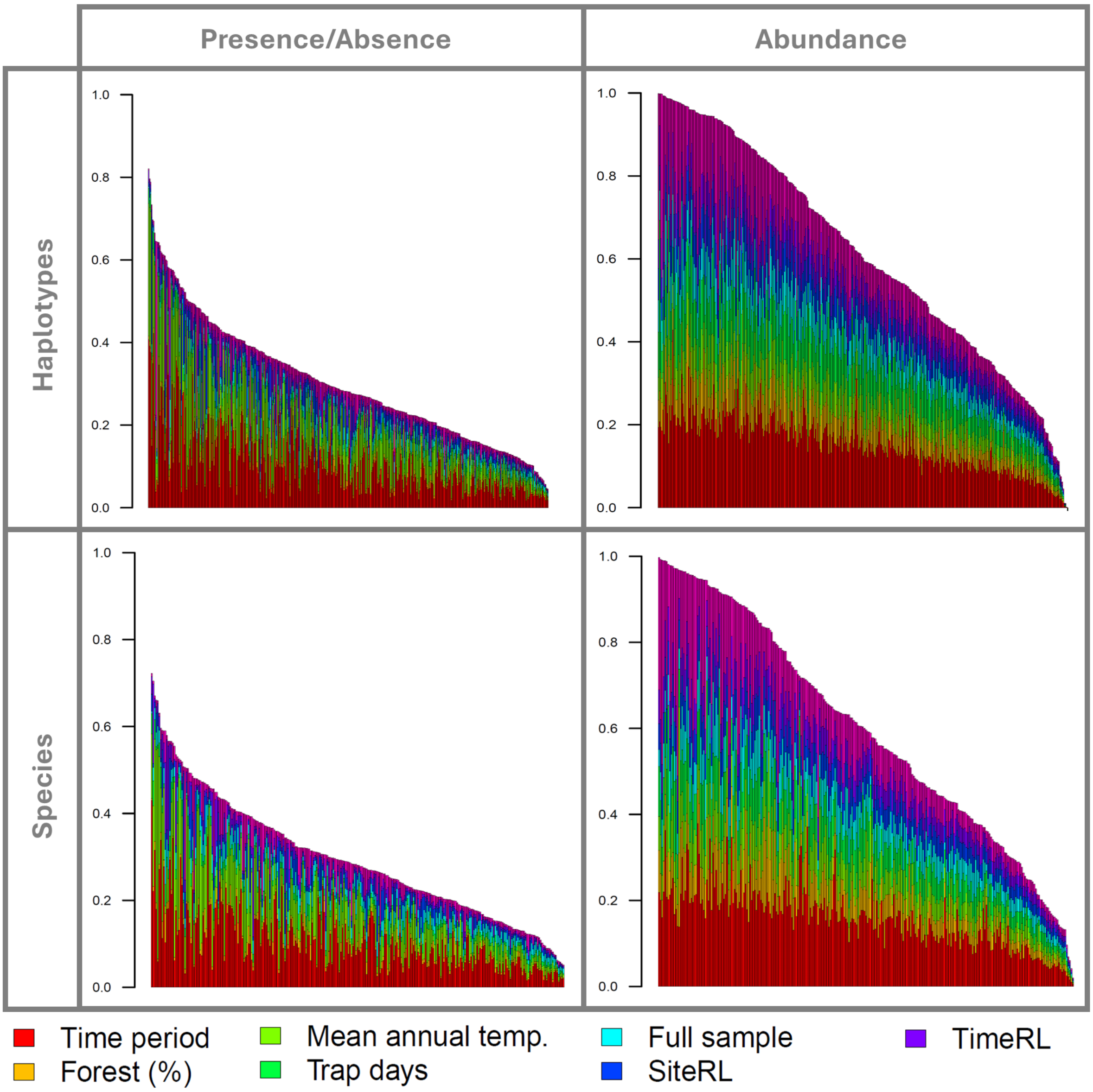


Supplementary Figure S8. Variance explained by fixed and random effects in HMSC models of the presence/absence or abundance of either haplotypes or species proxies.

*Each bar represents the variance partitioning result for a single haplotype or mOTU, illustrated in descending order of total explained variance (%).*

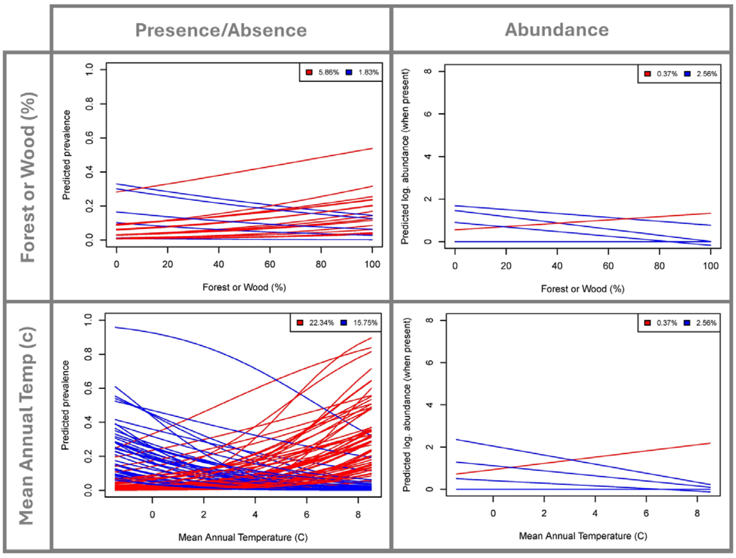


Supplementary Figure S9. Predicted mean prevalence and log(abundance) of species proxies as a function of forest or woodland cover (%) and mean annual temperature (°C) during late summer in HMSC models.

*Only those mOTUs are illustrated for which a directional trend was predicted with > 95% posterior support. The numbers in the legend represent the percentages of mOTUs, of the total modelled, that were predicted to have a positive (red) or negative (blue) response to the focal covariate. These are marginal predictions, where values for all covariates apart from the focal covariate and sampling season (late summer) were fixed at their mean value in the dataset. The results were similar for other sampling seasons as well, but only those for late summer are illustrated.*

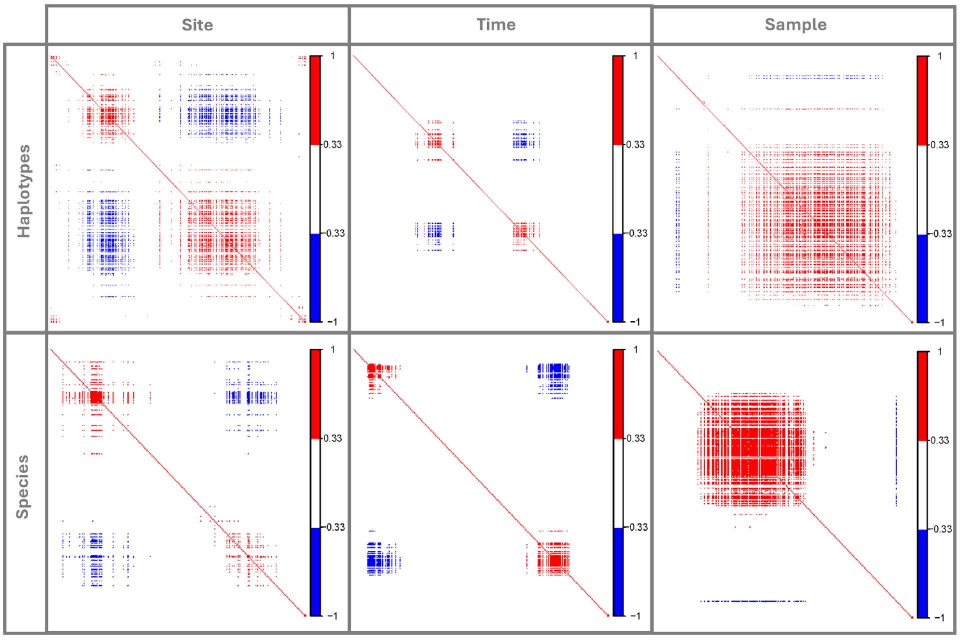


Supplementary Figure S10. Pairwise residual associations among haplotypes and species proxies in space and time as detected in HMSC models.

*Taxa are ordered on the x- and y-axes such that positive and negative residual associations are in blocks so that their prevalence across the datasets is easy to compare.*

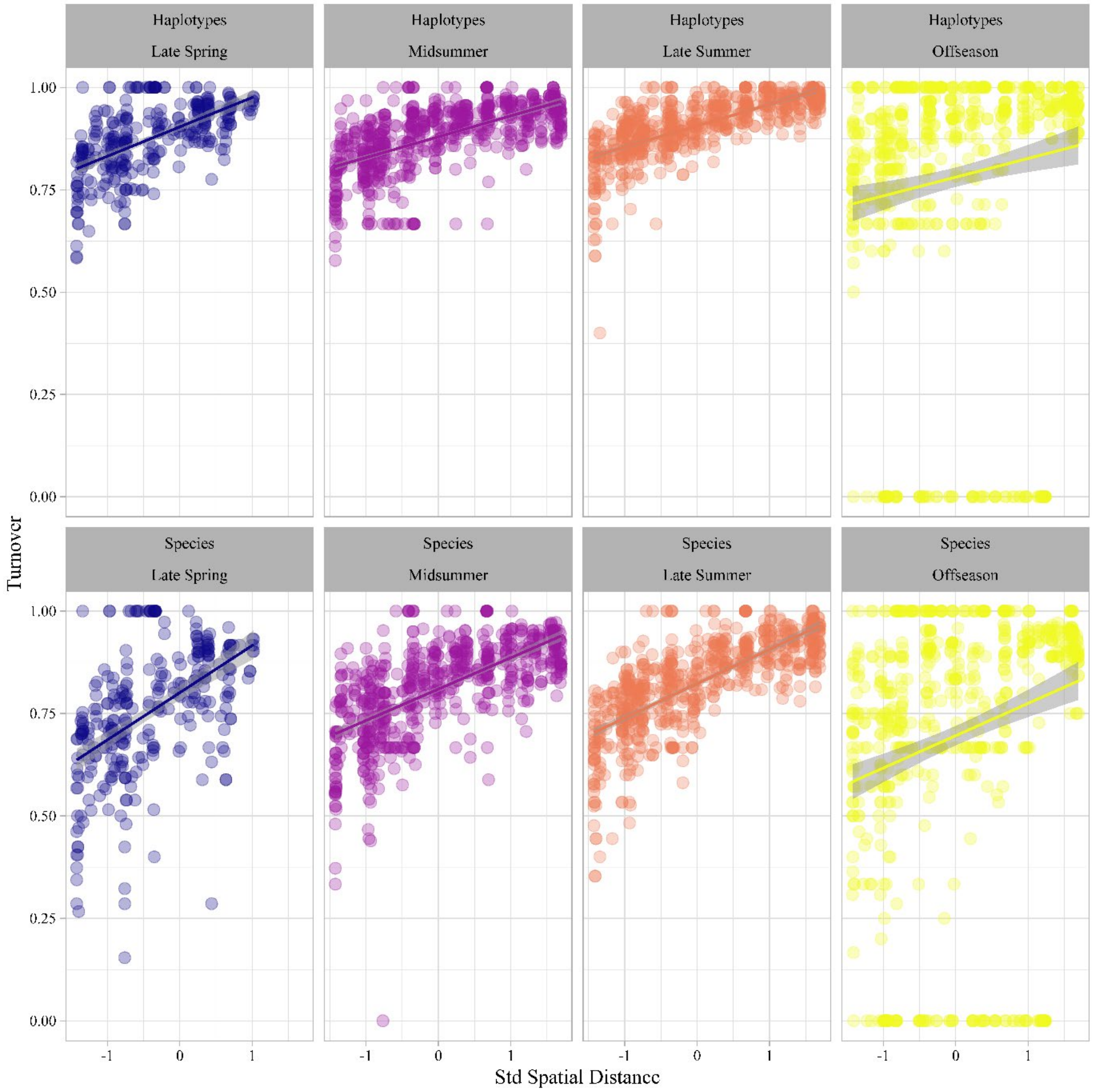


Supplementary Figure S11. Pairwise compositional turnover as a function of intervening geographic distance for each time period in the mOTU and Haplotype datasets.

*Following Baselga (2010), we partitioned overall Jaccard dissimilarity between samples from each time period (Late spring, Midsummer, Late summer, Offseason) into its turnover and nestedness components using either species (1.7% mOTUs) or haplotypes. Shown in each panel here are scatterplots of turnover of haplotypes (upper panels) vs mOTUs (lower panels) per time period between each pair of sampling sites as a function of their standardized intervening geographical distance.*

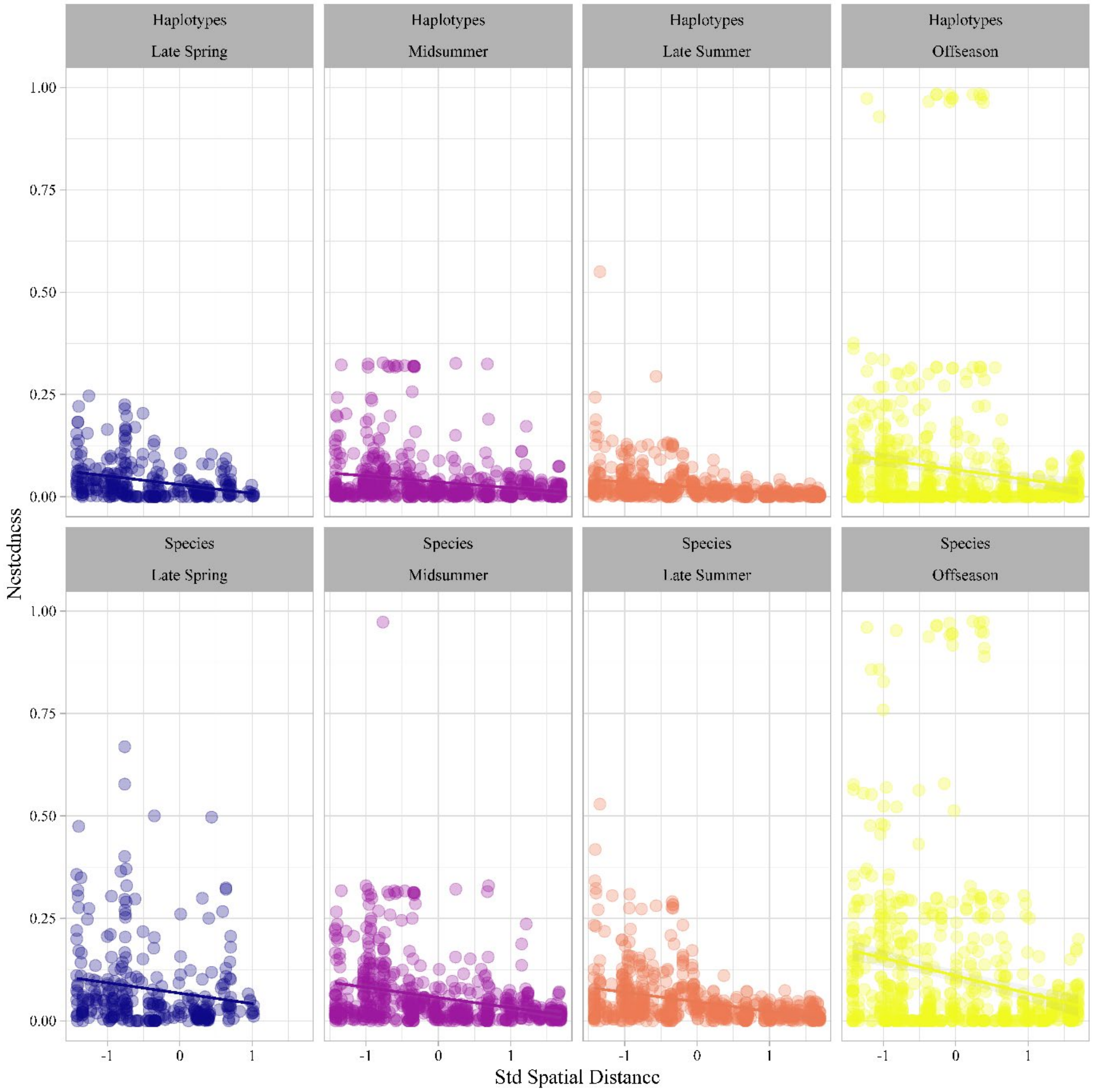


Supplementary Figure S12. Nestedness values scored along geographic distance gradients for each time period among the Species and Haplotype datasets.

*Following Baselga (2010), we partition overall Jaccard dissimilarity into its turnover and nestedness components using either species (1.7% mOTUs) or haplotypes recorded at each different time period. Shown in each panel are the scatterplots of nestedness between each pair of sampling sites (for each dataset) at each time period, as a function of their standardized intervening geographical distance.*

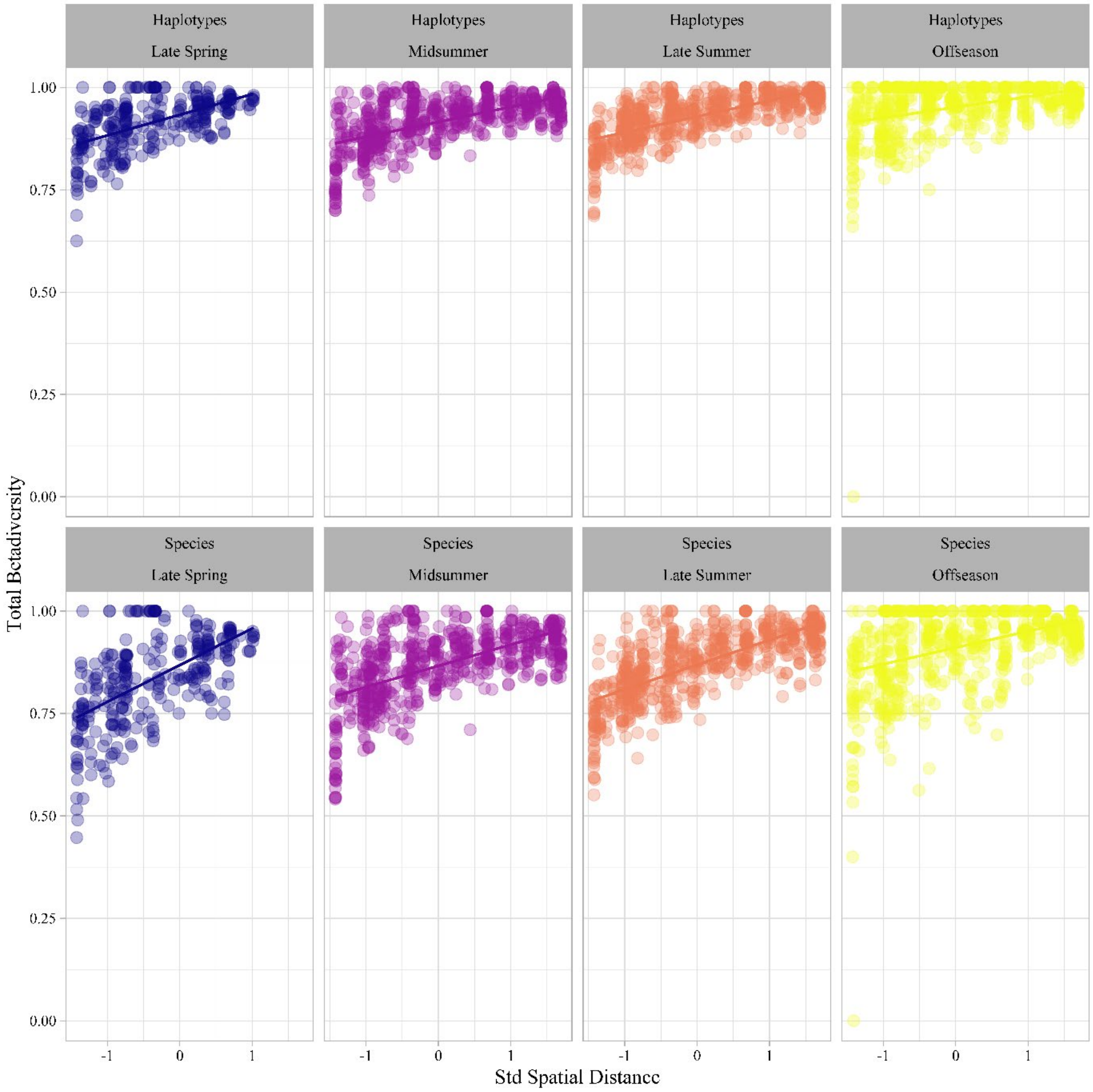


Supplementary Figure S13. Total beta-diversity values scored along geographic distance gradients for each time period among the Species and Haplotype datasets.

*Shown in each panel are the scatterplots of pairwise overall community dissimilarity between each pair of sampling sites (for each dataset) at each time period, against their spatial separation (standardized Euclidean distance between sample site coordinates). To illustrate the (lack of) impact of species delimitation criteria used, we show the same patterns using haplotypes (upper panels) vs species (1.7% mOTUs, lower panels).*

### Supplementary Table S1. Sample information.

*In this table, we provide metadata on each sample, grouped into the following columns: Site ID: the identity of the site; Start date: the first day of the sampling period (format day/month/year); End date: the first day of the sampling period (format day/month/year); Trapping days: length of sampling period (i.e. end date minus starting date); Latitude (in decimal degrees); Longitude (in decimal degrees); County (i.e., region within Sweden); Barcodes: the number of barcode sequences successfully retrieved; Sample sequence: information on whether all individuals in the sample were sequenced (Y) or not (N; see Supplementary Text S2); Time period: phase of summer phenology, grouped into: Late spring (May), midsummer (June), late summer (July/August), and offseason (the last sample of the season, collected in November or December 2018 or spring of 2019)(time periods are approximate, consult table for exact sample dates); Horticultural zone: plant hardiness zones defined by the Swedish Horticultural Society (Riksförbundet Svensk Trädgård 2018), with colours corresponding to those used in Fig. 1 of the main text); Avg annual temperature: mean annual temperature of the climate zone, derived from the Swedish Meteorological and Hydrological Institute (SMHI; smhi.se) for time period 1991-2020; Avg annual precipitation: mean annual precipitation of the climate zone, derived from the Swedish Meteorological and Hydrological Institute (SMHI; smhi.se) for time period 1991-2020.*

| Site ID | Sample ID | Start date | End date | Trapping Days | Latitude | Longitude | County | Barcodes | Entire sample sequenced | Time period | Horticultural zone | Avg. annual temperature | Avg. annual precipitation |
|---|---|---|---|---|---|---|---|---|---|---|---|---|---|
| 1 | M01013 | 12.10.2018 | 10.11.2018 | 29 | 55,52527 | 12,913264 | Skåne | 61 | Y | Offseason | I (red) | 8-10 | 500-600 |
| 1 | M01025 | 14.05.2018 | 28.05.2018 | 14 | 55,52527 | 12,913263 | Skåne | 312 | Y | Late Spring | I (red) | 8-10 | 500-600 |
| 1 | M01030 | 10.06.2018 | 25.06.2018 | 15 | 55,52527 | 12,913263 | Skåne | 214 | N | Midsummer | I (red) | 8-10 | 500-600 |
| 1 | M01035 | 19.07.2018 | 07.08.2018 | 19 | 55,52527 | 12,913263 | Skåne | 168 | N | Late Summer | I (red) | 8-10 | 500-600 |
| 2 | M02006 | 15.05.2018 | 25.05.2018 | 10 | 55,56939 | 12,926716 | Skåne | 535 | Y | Late Spring | I (red) | 8-10 | 500-600 |
| 2 | M02010 | 20.06.2018 | 28.06.2018 | 8 | 55,56939 | 12,926716 | Skåne | 184 | N | Midsummer | I (red) | 8-10 | 500-600 |
| 2 | M02014 | 30.07.2018 | 13.08.2018 | 14 | 55,56939 | 12,926716 | Skåne | 155 | N | Late Summer | I (red) | 8-10 | 500-600 |
| 2 | M02020 | 10.10.2018 | 05.11.2018 | 26 | 55,56939 | 12,926717 | Skåne | 35 | Y | Offseason | I (red) | 8-10 | 500-600 |
| 3 | M03006 | 15.05.2018 | 27.05.2018 | 12 | 55,63332 | 13,009706 | Skåne | 179 | Y | Late Spring | I (red) | 8-10 | 500-600 |
| 3 | M03010 | 20.06.2018 | 28.06.2018 | 8 | 55,63332 | 13,009706 | Skåne | 18 | Y | Midsummer | I (red) | 8-10 | 500-600 |
| 3 | M03014 | 30.07.2018 | 13.08.2018 | 14 | 55,63332 | 13,009706 | Skåne | 175 | N | Late Summer | I (red) | 8-10 | 500-600 |
| 3 | M03019 | 10.10.2018 | 05.11.2018 | 26 | 55,63332 | 13,009707 | Skåne | 111 | Y | Offseason | I (red) | 8-10 | 500-600 |
| 4 | M04006 | 15.05.2018 | 25.05.2018 | 10 | 55,57838 | 12,948558 | Skåne | 767 | N | Late Spring | I (red) | 8-10 | 500-600 |
| 4 | M04010 | 20.06.2018 | 28.06.2018 | 8 | 55,57838 | 12,948558 | Skåne | 261 | N | Midsummer | I (red) | 8-10 | 500-600 |
| 4 | M04014 | 30.07.2018 | 13.08.2018 | 14 | 55,57838 | 12,948558 | Skåne | 165 | N | Late Summer | I (red) | 8-10 | 500-600 |
| 4 | M04020 | 10.10.2018 | 05.11.2018 | 26 | 55,57838 | 12,948559 | Skåne | 182 | N | Offseason | I (red) | 8-10 | 500-600 |
| 5 | M05006 | 10.05.2018 | 19.05.2018 | 9 | 55,70427 | 13,452154 | Skåne | 525 | N | Late Spring | I (red) | 8-10 | 600-700 |
| 5 | M05012 | 16.06.2018 | 25.06.2018 | 9 | 55,70427 | 13,452154 | Skåne | 168 | N | Midsummer | I (red) | 8-10 | 600-700 |
| 5 | M05019 | 27.07.2018 | 07.08.2018 | 11 | 55,70427 | 13,452154 | Skåne | 172 | N | Late Summer | I (red) | 8-10 | 600-700 |
| 5 | M05029 | 30.10.2018 | 18.11.2018 | 19 | 55,70427 | 13,452155 | Skåne | 86 | Y | Offseason | I (red) | 8-10 | 600-700 |
| 6 | M06005 | 12.05.2018 | 19.05.2018 | 7 | 55,69799 | 13,496168 | Skåne | 421 | Y | Late Spring | I (red) | 8-10 | 600-700 |
| 6 | M06009 | 16.06.2018 | 25.06.2018 | 9 | 55,69799 | 13,496168 | Skåne | 178 | N | Midsummer | I (red) | 8-10 | 600-700 |
| 6 | M06016 | 27.07.2018 | 07.08.2018 | 11 | 55,69799 | 13,496168 | Skåne | 178 | N | Late Summer | I (red) | 8-10 | 600-700 |
| 6 | M06026 | 30.10.2018 | 18.11.2018 | 19 | 55,69799 | 13,496169 | Skåne | 24 | Y | Offseason | I (red) | 8-10 | 600-700 |
| 7 | M07004 | 14.05.2018 | 23.05.2018 | 9 | 55,77455 | 14,121564 | Skåne | 556 | N | Late Spring | I (red) | 8-10 | 700-800 |
| 7 | M07008 | 19.06.2018 | 28.06.2018 | 9 | 55,77455 | 14,121564 | Skåne | 181 | N | Midsummer | I (red) | 8-10 | 700-800 |
| 7 | M07012 | 26.07.2018 | 04.08.2018 | 9 | 55,77455 | 14,121564 | Skåne | 179 | N | Late Summer | I (red) | 8-10 | 700-800 |
| 7 | M07020 | 21.10.2018 | 06.11.2018 | 16 | 55,77455 | 14,121565 | Skåne | 182 | N | Offseason | I (red) | 8-10 | 700-800 |
| 8 | M08002 | 07.05.2018 | 21.05.2018 | 14 | 56,73076 | 13,067036 | Halland | 195 | Y | Late Spring | I (red) | 6-8 | 1100-1200 |
| 8 | M08005 | 12.06.2018 | 27.06.2018 | 15 | 56,73076 | 13,067036 | Halland | 161 | N | Midsummer | I (red) | 6-8 | 1100-1200 |
| 8 | M08009 | 29.07.2018 | 15.08.2018 | 17 | 56,73076 | 13,067036 | Halland | 115 | N | Late Summer | I (red) | 6-8 | 1100-1200 |
| 8 | M08013 | 29.10.2018 | 27.11.2018 | 29 | 56,73077 | 13,067037 | Halland | 9 | Y | Offseason | I (red) | 6-8 | 1100-1200 |
| 10 | M10001 | 18.05.2018 | 25.05.2018 | 7 | 56,22829 | 16,441877 | Kalmar | 482 | Y | Late Spring | I (red) | 8-10 | 400-500 |
| 10 | M10005 | 20.06.2018 | 29.06.2018 | 9 | 56,22829 | 16,441877 | Kalmar | 170 | N | Midsummer | I (red) | 8-10 | 400-500 |
| 10 | M10010 | 27.07.2018 | 03.08.2018 | 7 | 56,22829 | 16,441877 | Kalmar | 185 | N | Late Summer | I (red) | 8-10 | 400-500 |
| 10 | M10017 | 26.09.2018 | 21.12.2018 | 86 | 56,22829 | 16,441877 | Kalmar | 187 | N | Offseason | I (red) | 8-10 | 400-500 |
| 15 | M15002 | 15.06.2018 | 03.07.2018 | 18 | 58,33087 | 14,817465 | Östergötland | 313 | N | Midsummer | II (orange) | 6-8 | 500-600 |
| 15 | M15004 | 18.07.2018 | 06.08.2018 | 19 | 58,33087 | 14,817465 | Östergötland | 165 | N | Late Summer | II (orange) | 6-8 | 500-600 |
| 15 | M15010 | 14.10.2018 | 30.11.2018 | 47 | 58,33087 | 14,817466 | Östergötland | 134 | Y | Offseason | II (orange) | 6-8 | 500-600 |
| 15 | M15026 | 14.05.2018 | 24.05.2018 | 10 | 58,33087 | 14,817465 | Östergötland | 516 | N | Late Spring | II (orange) | 6-8 | 500-600 |
| 16 | M16002 | 14.05.2018 | 24.05.2018 | 10 | 58,33338 | 14,82316 | Östergötland | 547 | N | Late Spring | II (orange) | 6-8 | 500-600 |
| 16 | M16004 | 15.06.2018 | 03.07.2018 | 18 | 58,33338 | 14,82316 | Östergötland | 188 | N | Midsummer | II (orange) | 6-8 | 500-600 |
| 16 | M16008 | 18.07.2018 | 06.08.2018 | 19 | 58,33338 | 14,82316 | Östergötland | 156 | N | Late Summer | II (orange) | 6-8 | 500-600 |
| 16 | M16014 | 16.10.2018 | 30.11.2018 | 45 | 58,33338 | 14,82317 | Östergötland | 174 | N | Offseason | II (orange) | 6-8 | 500-600 |
| 17 | M17005 | 17.05.2018 | 23.05.2018 | 6 | 58,3442 | 11,334593 | Västra Götaland | 50 | Y | Late Spring | III (gold) | 8-10 | 800-900 |
| 17 | M17008 | 11.06.2018 | 22.06.2018 | 11 | 58,3442 | 11,334593 | Västra Götaland | 27 | Y | Midsummer | III (gold) | 8-10 | 800-900 |
| 17 | M17012 | 25.07.2018 | 09.08.2018 | 15 | 58,3442 | 11,334593 | Västra Götaland | 48 | Y | Late Summer | III (gold) | 8-10 | 800-900 |
| 17 | M17018 | 19.10.2018 | 05.02.2019 | 109 | 58,3442 | 11,334594 | Västra Götaland | 10 | Y | Offseason | III (gold) | 8-10 | 800-900 |
| 18 | M18005 | 13.05.2018 | 22.05.2018 | 9 | 59,27815 | 18,761432 | Stockholm | 1815 | Y | Late Spring | III (gold) | 6-8 | 500-600 |
| 18 | M18009 | 11.06.2018 | 22.06.2018 | 11 | 59,27815 | 18,761432 | Stockholm | 180 | N | Midsummer | III (gold) | 6-8 | 500-600 |
| 18 | M18015 | 29.07.2018 | 06.08.2018 | 8 | 59,27815 | 18,761432 | Stockholm | 176 | N | Late Summer | III (gold) | 6-8 | 500-600 |
| 18 | M18022 | 21.10.2018 | 16.11.2018 | 26 | 59,27815 | 18,761433 | Stockholm | 69 | Y | Offseason | III (gold) | 6-8 | 500-600 |
| 19 | M19007 | 17.05.2018 | 24.05.2018 | 7 | 59,33624 | 18,067797 | Stockholm | 1 | Y | Late Spring | II (orange) | 6-8 | 500-600 |
| 19 | M19012 | 21.06.2018 | 28.06.2018 | 7 | 59,33624 | 18,067797 | Stockholm | 2 | Y | Midsummer | II (orange) | 6-8 | 500-600 |
| 19 | M19017 | 25.07.2018 | 02.08.2018 | 8 | 59,33624 | 18,067797 | Stockholm | 11 | Y | Late Summer | II (orange) | 6-8 | 500-600 |
| 19 | M19027 | 05.10.2018 | 01.11.2018 | 27 | 59,33624 | 18,067798 | Stockholm | 1 | Y | Offseason | II (orange) | 6-8 | 500-600 |
| 20 | M20002 | 16.05.2018 | 28.05.2018 | 12 | 62,44088 | 17,423142 | Medelpad | 562 | N | Late Spring | IV (olive) | 4-6 | 500-600 |
| 20 | M20005 | 13.06.2018 | 23.06.2018 | 10 | 62,44088 | 17,423142 | Medelpad | 188 | N | Midsummer | IV (olive) | 4-6 | 500-600 |
| 20 | M20011 | 23.07.2018 | 01.08.2018 | 9 | 62,44088 | 17,423142 | Medelpad | 181 | N | Late Summer | IV (olive) | 4-6 | 500-600 |
| 20 | M20020 | 29.10.2018 | 15.11.2018 | 17 | 62,44088 | 17,423142 | Medelpad | 35 | Y | Offseason | IV (olive) | 4-6 | 500-600 |
| 21 | M21002 | 15.05.2018 | 29.05.2018 | 14 | 63,34501 | 14,537859 | Jämtland | 551 | N | Late Spring | VI (purple) | 2-4 | 500-600 |
| 21 | M21004 | 12.06.2018 | 25.06.2018 | 13 | 63,34501 | 14,537859 | Jämtland | 260 | N | Midsummer | VI (purple) | 2-4 | 500-600 |
| 21 | M21007 | 23.07.2018 | 06.08.2018 | 14 | 63,34501 | 14,537859 | Jämtland | 179 | N | Late Summer | VI (purple) | 2-4 | 500-600 |
| 21 | M21014 | 30.10.2018 | 12.11.2018 | 13 | 63,34501 | 14,537859 | Jämtland | 2 | Y | Offseason | VI (purple) | 2-4 | 500-600 |
| 22 | M22003 | 17.05.2018 | 24.05.2018 | 7 | 62,6455 | 15,63441 | Jämtland | 563 | N | Late Spring | VII (dark blue) | 2-4 | 500-600 |
| 22 | M22006 | 11.06.2018 | 22.06.2018 | 11 | 62,6455 | 15,63441 | Jämtland | 180 | N | Midsummer | VII (dark blue) | 2-4 | 500-600 |

| | | | | | | | | | | | | | |
|---|---|---|---|---|---|---|---|---|---|---|---|---|---|
| 22 | M22012 | 28.07.2018 | 05.08.2018 | 8 | 62,6455 | 15,63441 | Jämtland | 172 | N | Late Summer | VII (dark blue) | 2-4 | 500-600 |
| 22 | M22018 | 14.10.2018 | 04.11.2018 | 21 | 62,6455 | 15,63441 | Jämtland | 94 | Y | Offseason | VII (dark blue) | 2-4 | 500-600 |
| 23 | M23001 | 16.05.2018 | 23.05.2018 | 7 | 63,8201 | 20,316182 | Västerbotten | 301 | Y | Late Spring | V (green) | 4-6 | 600-700 |
| 23 | M23006 | 20.06.2018 | 26.06.2018 | 6 | 63,8201 | 20,316182 | Västerbotten | 179 | N | Midsummer | V (green) | 4-6 | 600-700 |
| 23 | M23011 | 25.07.2018 | 01.08.2018 | 7 | 63,8201 | 20,316182 | Västerbotten | 186 | N | Late Summer | V (green) | 4-6 | 600-700 |
| 23 | M23014 | 26.09.2018 | 07.11.2018 | 42 | 63,8201 | 20,316182 | Västerbotten | 55 | Y | Offseason | V (green) | 4-6 | 600-700 |
| 24 | M24002 | 20.05.2018 | 28.05.2018 | 8 | 63,79557 | 20,897138 | Västerbotten | 288 | Y | Late Spring | V (green) | 4-6 | 500-600 |
| 24 | M24006 | 18.06.2018 | 24.06.2018 | 6 | 63,79557 | 20,897138 | Västerbotten | 189 | N | Midsummer | V (green) | 4-6 | 500-600 |
| 24 | M24013 | 26.07.2018 | 05.08.2018 | 10 | 63,79557 | 20,897138 | Västerbotten | 170 | N | Late Summer | V (green) | 4-6 | 500-600 |
| 24 | M24018 | 11.10.2018 | 14.11.2018 | 34 | 63,79557 | 20,897138 | Västerbotten | 37 | Y | Offseason | V (green) | 4-6 | 500-600 |
| 25 | M25001 | 01.06.2018 | 24.06.2018 | 23 | 66,42998 | 20,624348 | Norrbotten | 542 | N | Midsummer | VI (purple) | 0-2 | 600-700 |
| 25 | M25004 | 22.07.2018 | 07.08.2018 | 16 | 66,42998 | 20,624348 | Norrbotten | 172 | N | Late Summer | VI (purple) | 0-2 | 600-700 |
| 25 | M25007 | 25.10.2018 | 15.11.2018 | 21 | 66,42998 | 20,624349 | Norrbotten | no phorids | n/a | Offseason | VI (purple) | 0-2 | 600-700 |
| 26 | M26001 | 02.06.2018 | 24.06.2018 | 22 | 66,37515 | 20,713693 | Norrbotten | 545 | N | Midsummer | VI (purple) | 0-2 | 600-700 |
| 26 | M26004 | 22.07.2018 | 07.08.2018 | 16 | 66,37515 | 20,713693 | Norrbotten | 183 | N | Late Summer | VI (purple) | 0-2 | 600-700 |
| 26 | M26007 | 25.10.2018 | 15.11.2018 | 21 | 66,37515 | 20,713694 | Norrbotten | no phorids | n/a | Offseason | VI (purple) | 0-2 | 600-700 |
| 28 | M28001 | 05.06.2018 | 28.06.2018 | 23 | 68,11062 | 23,327608 | Norrbotten | 545 | N | Midsummer | VIII (light blue) | -2-0 | 500-600 |
| 28 | M28007 | 25.07.2018 | 01.08.2018 | 7 | 68,11062 | 23,327608 | Norrbotten | 183 | N | Late Summer | VIII (light blue) | -2-0 | 500-600 |
| 28 | M28010 | 29.09.2018 | 25.05.2019 | 238 | 68,11062 | 23,327608 | Norrbotten | 184 | N | Offseason | VIII (light blue) | -2-0 | 500-600 |
| 29 | M29001 | 05.06.2018 | 28.06.2018 | 23 | 68,07975 | 23,243206 | Norrbotten | 500 | N | Midsummer | VIII (light blue) | -2-0 | 500-600 |
| 29 | M29007 | 25.07.2018 | 03.08.2018 | 9 | 68,07975 | 23,243206 | Norrbotten | 185 | N | Late Summer | VIII (light blue) | -2-0 | 500-600 |
| 29 | M29009 | 18.08.2018 | 07.07.2019 | 323 | 68,07975 | 23,243206 | Norrbotten | 186 | N | Offseason | VIII (light blue) | -2-0 | 500-600 |
| 30 | M30001 | 05.06.2018 | 28.06.2018 | 23 | 68,11143 | 23,330286 | Norrbotten | 174 | N | Midsummer | VIII (light blue) | -2-0 | 500-600 |
| 30 | M30007 | 25.07.2018 | 01.08.2018 | 7 | 68,11143 | 23,330286 | Norrbotten | 188 | N | Late Summer | VIII (light blue) | -2-0 | 500-600 |
| 30 | M30010 | 29.09.2018 | 02.07.2019 | 276 | 68,11143 | 23,330286 | Norrbotten | 187 | N | Offseason | VIII (light blue) | -2-0 | 500-600 |
| 31 | M31001 | 03.06.2018 | 03.07.2018 | 30 | 68,35478 | 18,822478 | Norrbotten | 503 | N | Midsummer | Alpine (white) | 0-2 | 300-400 |
| 31 | M31005 | 29.07.2018 | 07.08.2018 | 9 | 68,35478 | 18,822478 | Norrbotten | 187 | N | Late Summer | Alpine (white) | 0-2 | 300-400 |
| 31 | M31013 | 04.10.2018 | 27.05.2019 | 235 | 68,35478 | 18,822478 | Norrbotten | 186 | N | Offseason | Alpine (white) | 0-2 | 300-400 |
| 32 | M32001 | 03.06.2018 | 03.07.2018 | 30 | 68,35515 | 18,826526 | Norrbotten | 531 | N | Midsummer | Alpine (white) | 0-2 | 300-400 |
| 32 | M32005 | 29.07.2018 | 07.08.2018 | 9 | 68,35515 | 18,826526 | Norrbotten | 178 | N | Late Summer | Alpine (white) | 0-2 | 300-400 |
| 32 | M32013 | 04.10.2018 | 27.05.2019 | 235 | 68,35515 | 18,826526 | Norrbotten | 189 | N | Offseason | Alpine (white) | 0-2 | 300-400 |
| 33 | M33001 | 03.06.2018 | 03.07.2018 | 30 | 68,36345 | 18,763011 | Norrbotten | 523 | N | Midsummer | Alpine (white) | 0-2 | 300-400 |
| 33 | M33005 | 29.07.2018 | 07.08.2018 | 9 | 68,36345 | 18,763011 | Norrbotten | 183 | N | Late Summer | Alpine (white) | 0-2 | 300-400 |
| 33 | M33013 | 04.10.2018 | 27.05.2019 | 235 | 68,36345 | 18,763011 | Norrbotten | 187 | N | Offseason | Alpine (white) | 0-2 | 300-400 |
| 34 | M34001 | 03.06.2018 | 03.07.2018 | 30 | 68,35441 | 18,827631 | Norrbotten | 542 | N | Midsummer | Alpine (white) | 0-2 | 300-400 |
| 34 | M34005 | 29.07.2018 | 07.08.2018 | 9 | 68,35441 | 18,827631 | Norrbotten | 180 | N | Late Summer | Alpine (white) | 0-2 | 300-400 |
| 34 | M34013 | 04.10.2018 | 28.05.2019 | 236 | 68,35441 | 18,827631 | Norrbotten | 187 | N | Offseason | Alpine (white) | 0-2 | 300-400 |
| 35 | M35001 | 03.06.2018 | 03.07.2018 | 30 | 68,36101 | 18,736851 | Norrbotten | 554 | N | Midsummer | Alpine (white) | 0-2 | 300-400 |
| 35 | M35005 | 29.07.2018 | 07.08.2018 | 9 | 68,36101 | 18,736851 | Norrbotten | 176 | N | Late Summer | Alpine (white) | 0-2 | 300-400 |
| 35 | M35013 | 04.10.2018 | 16.06.2019 | 255 | 68,36101 | 18,736851 | Norrbotten | 188 | N | Offseason | Alpine (white) | 0-2 | 300-400 |
| 38 | M38001 | 01.06.2018 | 15.06.2018 | 14 | 65,95729 | 16,205235 | Västerbotten | 525 | N | Late Spring | VIII (light blue) | 0-2 | 600-700 |
| 38 | M38002 | 15.06.2018 | 02.07.2018 | 17 | 65,95729 | 16,205235 | Västerbotten | 142 | N | Midsummer | VIII (light blue) | 0-2 | 600-700 |
| 38 | M38004 | 22.07.2018 | 17.08.2018 | 26 | 65,95729 | 16,205235 | Västerbotten | 183 | N | Late Summer | VIII (light blue) | 0-2 | 600-700 |
| 38 | M38008 | 19.10.2018 | 09.11.2018 | 21 | 65,95729 | 16,205235 | Västerbotten | 6 | Y | Offseason | VIII (light blue) | 0-2 | 600-700 |
| 40 | M40013 | 10.06.2018 | 25.06.2018 | 15 | 55,52346 | 12,910872 | Skåne | 166 | N | Midsummer | I (red) | 8-10 | 500-600 |
| 40 | M40014 | 14.05.2018 | 28.05.2018 | 14 | 55,52346 | 12,910872 | Skåne | 218 | N | Late Spring | I (red) | 8-10 | 500-600 |
| 40 | M40017 | 19.07.2018 | 07.08.2018 | 19 | 55,52346 | 12,910872 | Skåne | 155 | N | Late Summer | I (red) | 8-10 | 500-600 |
| 40 | M40023 | 12.10.2018 | 10.11.2018 | 29 | 55,52347 | 12,910873 | Skåne | 5 | Y | Offseason | I (red) | 8-10 | 500-600 |
| 41 | M41006 | 14.05.2018 | 22.05.2018 | 8 | 57,68907 | 11,956801 | Västergötland | 443 | Y | Late Spring | II (orange) | 8-10 | 900-1000 |
| 41 | M41011 | 19.06.2018 | 26.06.2018 | 7 | 57,68907 | 11,956801 | Västergötland | 156 | N | Midsummer | II (orange) | 8-10 | 900-1000 |
| 41 | M41016 | 26.07.2018 | 02.08.2018 | 7 | 57,68907 | 11,956801 | Västergötland | 180 | N | Late Summer | II (orange) | 8-10 | 900-1000 |
| 41 | M41022 | 12.10.2018 | 05.11.2018 | 24 | 57,68907 | 11,956801 | Västergötland | 171 | N | Offseason | II (orange) | 8-10 | 900-1000 |
| 42 | M42003 | 12.10.2018 | 10.11.2018 | 29 | 55,52382 | 12,911576 | Skåne | 15 | Y | Offseason | I (red) | 8-10 | 500-600 |
| 42 | M42004 | 19.07.2018 | 07.08.2018 | 19 | 55,52382 | 12,911575 | Skåne | 170 | N | Late Summer | I (red) | 8-10 | 500-600 |
| 42 | M42025 | 14.05.2018 | 28.05.2018 | 14 | 55,52382 | 12,911575 | Skåne | 636 | N | Late Spring | I (red) | 8-10 | 500-600 |
| 42 | M42030 | 10.06.2018 | 25.06.2018 | 15 | 55,52382 | 12,911575 | Skåne | 174 | N | Midsummer | I (red) | 8-10 | 500-600 |
| 43 | M43001 | 05.06.2018 | 28.06.2018 | 23 | 68,11132 | 23,331281 | Norrbotten | 557 | N | Midsummer | VIII (light blue) | -2-0 | 500-600 |
| 43 | M43007 | 25.07.2018 | 01.08.2018 | 7 | 68,11132 | 23,331281 | Norrbotten | 185 | N | Late Summer | VIII (light blue) | -2-0 | 500-600 |
| 43 | M43010 | 29.09.2018 | 20.07.2019 | 294 | 68,11132 | 23,331281 | Norrbotten | 187 | N | Offseason | VIII (light blue) | -2-0 | 500-600 |
| 44 | M44002 | 15.05.2018 | 29.05.2018 | 14 | 63,34467 | 14,536979 | Jämtland | 524 | N | Late Spring | VI (purple) | 2-4 | 500-600 |
| 44 | M44004 | 12.06.2018 | 25.06.2018 | 13 | 63,34467 | 14,536979 | Jämtland | 169 | N | Midsummer | VI (purple) | 2-4 | 500-600 |
| 44 | M44007 | 23.07.2018 | 06.08.2018 | 14 | 63,34467 | 14,536979 | Jämtland | 184 | N | Late Summer | VI (purple) | 2-4 | 500-600 |
| 44 | M44014 | 30.10.2018 | 12.11.2018 | 13 | 63,34467 | 14,536979 | Jämtland | 1 | Y | Offseason | VI (purple) | 2-4 | 500-600 |
| 45 | M45002 | 20.05.2018 | 28.05.2018 | 8 | 63,79621 | 20,900016 | Västerbotten | 63 | Y | Late Spring | V (green) | 4-6 | 500-600 |
| 45 | M45006 | 18.06.2018 | 24.06.2018 | 6 | 63,79621 | 20,900016 | Västerbotten | 64 | Y | Midsummer | V (green) | 4-6 | 500-600 |
| 45 | M45013 | 26.07.2018 | 05.08.2018 | 10 | 63,79621 | 20,900016 | Västerbotten | 179 | N | Late Summer | V (green) | 4-6 | 500-600 |
| 45 | M45018 | 11.10.2018 | 14.11.2018 | 34 | 63,79621 | 20,900016 | Västerbotten | 24 | Y | Offseason | V (green) | 4-6 | 500-600 |
| 46 | M46001 | 04.07.2018 | 12.07.2018 | 8 | 56,61953 | 16,498101 | Kalmar | 932 | N | Midsummer | I (red) | 6-8 | 500-600 |
| 46 | M46004 | 31.07.2018 | 30.08.2018 | 30 | 56,61953 | 16,498101 | Kalmar | 178 | N | Late Summer | I (red) | 6-8 | 500-600 |
| 46 | M46006 | 15.10.2018 | 01.01.2019 | 78 | 56,61953 | 16,498101 | Kalmar | 48 | Y | Offseason | I (red) | 6-8 | 500-600 |

### Supplementary Table S2. Distinctness of scuttle fly communities across zones, as based on different clustering thresholds for species delimitation.

*Species turnover across zones, assessed using ANOSIM. Values above the grey diagonal line are R-statistics, whereas values below the diagonal line are P-values (Significance codes: 0 '***' 0.001 '**' 0.01 '*' 0.05 '.' 0.1 ' ' 1) for haplotypes and mOTUs delimited at thresholds of 0.5%, 1.7%, 3%, 4%, and 5%.*

**ANOSIM(Haplo)** R statistic above diagonal — Significance level below diagonal
Sample statistic: 0.539 — Significance of sample statistic: 0.001 — Number of permutations: 999

| | Zone 1 | Zone 2 | Zone 3 | Zone 4 | Zone 5 | Zone 6 | Zone 7 | Zone 8 | Alpine zone |
|---|---|---|---|---|---|---|---|---|---|
| Zone 1 | | 0,309 | 0,189 | 0,481 | 0,662 | 0,668 | 0,739 | 0,783 | 0,899 |
| Zone 2 | 0,002 | | 0,233 | 0,312 | 0,661 | 0,588 | 0,705 | 0,567 | 0,626 |
| Zone 3 | 0,132 | 0,062 | | 0,593 | 0,571 | 0,546 | 0,778 | 0,454 | 0,289 |
| Zone 4 | 0,006 | 0,018 | 0,100 | | 0,369 | 0,142 | 0,185 | 0,331 | 0,252 |
| Zone 5 | 0,001 | 0,001 | 0,017 | 0,025 | | 0,227 | 0,321 | 0,228 | 0,238 |
| Zone 6 | 0,001 | 0,001 | 0,010 | 0,171 | 0,029 | | 0,193 | 0,145 | 0,256 |
| Zone 7 | 0,002 | 0,002 | 0,100 | 0,300 | 0,033 | 0,112 | | 0,248 | 0,214 |
| Zone 8 | 0,001 | 0,001 | 0,012 | 0,042 | 0,030 | 0,039 | 0,074 | | 0,156 |
| Alpine zone | 0,001 | 0,001 | 0,039 | 0,049 | 0,023 | 0,007 | 0,072 | 0,023 | |

**ANOSIM(0.5%)** R statistic above diagonal — Significance level below diagonal
Sample statistic: 0.539 — Significance of sample statistic: 0.001 — Number of permutations: 999

| | Zone 1 | Zone 2 | Zone 3 | Zone 4 | Zone 5 | Zone 6 | Zone 7 | Zone 8 | Alpine zone |
|---|---|---|---|---|---|---|---|---|---|
| Zone 1 | | 0,334 | 0,064 | 0,488 | 0,692 | 0,695 | 0,774 | 0,796 | 0,870 |
| Zone 2 | 0,001 | | 0,196 | 0,344 | 0,654 | 0,593 | 0,751 | 0,592 | 0,544 |
| Zone 3 | 0,323 | 0,110 | | 0,444 | 0,536 | 0,538 | 0,630 | 0,480 | 0,290 |
| Zone 4 | 0,004 | 0,024 | 0,200 | | 0,294 | 0,143 | 0,111 | 0,346 | 0,181 |
| Zone 5 | 0,001 | 0,001 | 0,017 | 0,058 | | 0,212 | 0,278 | 0,243 | 0,236 |
| Zone 6 | 0,001 | 0,001 | 0,010 | 0,161 | 0,033 | | 0,208 | 0,151 | 0,196 |
| Zone 7 | 0,001 | 0,002 | 0,100 | 0,500 | 0,075 | 0,105 | | 0,272 | 0,145 |
| Zone 8 | 0,001 | 0,001 | 0,007 | 0,032 | 0,014 | 0,033 | 0,049 | | 0,101 |
| Alpine zone | 0,001 | 0,001 | 0,036 | 0,097 | 0,017 | 0,021 | 0,118 | 0,052 | |

**ANOSIM(1.7%)** R statistic above diagonal — Significance level below diagonal
Sample statistic: 0.512 — Significance of sample statistic: 0.001 — Number of permutations: 999

| | Zone 1 | Zone 2 | Zone 3 | Zone 4 | Zone 5 | Zone 6 | Zone 7 | Zone 8 | Alpine zone |
|---|---|---|---|---|---|---|---|---|---|
| Zone 1 | | 0,326 | 0,074 | 0,472 | 0,643 | 0,640 | 0,716 | 0,765 | 0,877 |
| Zone 2 | 0,002 | | 0,217 | 0,297 | 0,675 | 0,517 | 0,757 | 0,602 | 0,601 |
| Zone 3 | 0,291 | 0,099 | | 0,481 | 0,579 | 0,486 | 0,556 | 0,492 | 0,461 |
| Zone 4 | 0,016 | 0,031 | 0,200 | | 0,298 | 0,049 | 0,111 | 0,287 | 0,205 |
| Zone 5 | 0,001 | 0,001 | 0,017 | 0,067 | | 0,156 | 0,286 | 0,199 | 0,226 |
| Zone 6 | 0,001 | 0,001 | 0,010 | 0,315 | 0,076 | | 0,149 | 0,153 | 0,223 |
| Zone 7 | 0,001 | 0,002 | 0,100 | 0,600 | 0,058 | 0,154 | | 0,223 | 0,173 |
| Zone 8 | 0,001 | 0,001 | 0,006 | 0,050 | 0,032 | 0,041 | 0,071 | | 0,095 |
| Alpine zone | 0,001 | 0,001 | 0,013 | 0,086 | 0,024 | 0,020 | 0,105 | 0,063 | |

**ANOSIM(3%)** R statistic above diagonal — Significance level below diagonal
Sample statistic: 0.466 — Significance of sample statistic: 0.001 — Number of permutations: 999

| | Zone 1 | Zone 2 | Zone 3 | Zone 4 | Zone 5 | Zone 6 | Zone 7 | Zone 8 | Alpine zone |
|---|---|---|---|---|---|---|---|---|---|
| Zone 1 | | 0,279 | 0,085 | 0,415 | 0,522 | 0,579 | 0,726 | 0,612 | 0,670 |
| Zone 2 | 0,002 | | 0,273 | 0,244 | 0,575 | 0,526 | 0,782 | 0,580 | 0,749 |
| Zone 3 | 0,289 | 0,062 | | 0,481 | 0,516 | 0,431 | 0,519 | 0,556 | 0,931 |
| Zone 4 | 0,060 | 0,035 | 0,200 | | 0,286 | 0,017 | 0,000 | 0,311 | 0,867 |
| Zone 5 | 0,001 | 0,001 | 0,025 | 0,042 | | 0,111 | 0,385 | 0,281 | 0,766 |
| Zone 6 | 0,001 | 0,001 | 0,010 | 0,399 | 0,137 | | 0,108 | 0,195 | 0,574 |
| Zone 7 | 0,001 | 0,002 | 0,100 | 0,600 | 0,033 | 0,199 | | 0,351 | 0,793 |
| Zone 8 | 0,001 | 0,001 | 0,001 | 0,025 | 0,015 | 0,013 | 0,017 | | 0,248 |
| Alpine zone | 0,001 | 0,001 | 0,001 | 0,001 | 0,001 | 0,001 | 0,001 | 0,002 | |

**ANOSIM(4%)** R statistic above diagonal Significance level below diagonal

Sample statistic: 0.399 Significance of sample statistic: 0.001 Number of permutations: 999

| | Zone 1 | Zone 2 | Zone 3 | Zone 4 | Zone 5 | Zone 6 | Zone 7 | Zone 8 | Alpine zone |
|---|---|---|---|---|---|---|---|---|---|
| Zone 1 | | 0,267 | 0,101 | 0,419 | 0,487 | 0,548 | 0,686 | 0,523 | 0,540 |
| Zone 2 | 0,002 | | 0,260 | 0,194 | 0,391 | 0,452 | 0,713 | 0,510 | 0,634 |
| Zone 3 | 0,291 | 0,099 | | 0,556 | 0,496 | 0,506 | 0,630 | 0,661 | 0,972 |
| Zone 4 | 0,016 | 0,031 | 0,200 | | 0,254 | 0,014 | 0,111 | 0,242 | 0,843 |
| Zone 5 | 0,001 | 0,001 | 0,017 | 0,067 | | 0,082 | 0,337 | 0,242 | 0,693 |
| Zone 6 | 0,001 | 0,001 | 0,010 | 0,315 | 0,076 | | 0,129 | 0,203 | 0,620 |
| Zone 7 | 0,001 | 0,002 | 0,100 | 0,600 | 0,058 | 0,154 | | 0,299 | 0,809 |
| Zone 8 | 0,001 | 0,001 | 0,006 | 0,050 | 0,032 | 0,041 | 0,071 | | 0,271 |
| Alpine zone | 0,001 | 0,001 | 0,013 | 0,086 | 0,024 | 0,020 | 0,105 | 0,063 | |

**ANOSIM(5%)** R statistic above diagonal Significance level below diagonal

Sample statistic: 0.270 Significance of sample statistic: 0.001 Number of permutations: 999

| | Zone 1 | Zone 2 | Zone 3 | Zone 4 | Zone 5 | Zone 6 | Zone 7 | Zone 8 | Alpine zone |
|---|---|---|---|---|---|---|---|---|---|
| Zone 1 | | 0,181 | 0,228 | 0,366 | 0,468 | 0,408 | 0,447 | 0,408 | 0,319 |
| Zone 2 | 0,006 | | 0,425 | 0,267 | 0,332 | 0,317 | 0,513 | 0,351 | 0,444 |
| Zone 3 | 0,700 | 0,022 | | 0,593 | 0,540 | 0,601 | 0,519 | 0,740 | 0,955 |
| Zone 4 | 0,002 | 0,044 | 0,100 | | 0,119 | 0,053 | -0,037 | 0,238 | 0,799 |
| Zone 5 | 0,001 | 0,003 | 0,008 | 0,242 | | 0,159 | 0,234 | 0,285 | 0,676 |
| Zone 6 | 0,001 | 0,003 | 0,003 | 0,301 | 0,063 | | 0,154 | 0,212 | 0,584 |
| Zone 7 | 0,001 | 0,002 | 0,100 | 0,600 | 0,108 | 0,143 | | 0,373 | 0,785 |
| Zone 8 | 0,001 | 0,001 | 0,001 | 0,070 | 0,006 | 0,100 | 0,009 | | 0,256 |
| Alpine zone | 0,001 | 0,001 | 0,001 | 0,002 | 0,001 | 0,001 | 0,002 | 0,002 | |

### Supplementary Table S3. Similarity of scuttle fly communities across zones, as based on different clustering thresholds for species delimitation.

*Shown are results of SIMPER analyses of species turnover across zones, for haplotypes and mOTUs delimited at thresholds of 0.5%, 1.7%, 3%, 4%, and 5%.*

**SIMPER(Haplo)**

| | Within Zone Similarity (%) | Between Zone Similarity (%) Zone 1 | Zone 2 | Zone 3 | Zone 4 | Zone 5 | Zone 6 | Zone 7 | Zone 8 | Alpine zone |
|---|---|---|---|---|---|---|---|---|---|---|
| Zone 1 | 18,32 | | | | | | | | | |
| Zone 2 | 18,75 | 14,03 | | | | | | | | |
| Zone 3 | 22,81 | 15,32 | 15,15 | | | | | | | |
| Zone 4 | 23,29 | 11,88 | 14,06 | 15,83 | | | | | | |
| Zone 5 | 21,5 | 9,75 | 9,78 | 12,94 | 16,28 | | | | | |
| Zone 6 | 22,19 | 9,51 | 11,46 | 12,85 | 18,89 | 17,76 | | | | |
| Zone 7 | 18,78 | 8,45 | 8,67 | 11,87 | 19,48 | 16,20 | 18,38 | | | |
| Zone 8 | 18,67 | 6,94 | 8,50 | 9,08 | 11,30 | 14,15 | 16,50 | 13,03 | | |
| Alpine zone | 17,75 | 3,27 | 3,74 | 4,95 | 5,98 | 8,15 | 10,20 | 8,14 | 13,24 | |

**SIMPER(0.5%)**

| | Within Zone Similarity (%) | Between Zone Similarity (%) Zone 1 | Zone 2 | Zone 3 | Zone 4 | Zone 5 | Zone 6 | Zone 7 | Zone 8 | Alpine zone |
|---|---|---|---|---|---|---|---|---|---|---|
| Zone 1 | 24,3 | | | | | | | | | |
| Zone 2 | 24,41 | 18,83 | | | | | | | | |
| Zone 3 | 29,45 | 22,63 | 20,8 | | | | | | | |
| Zone 4 | 28,75 | 16,19 | 18,56 | 21,81 | | | | | | |
| Zone 5 | 26,03 | 13,16 | 13,42 | 16,87 | 21,41 | | | | | |
| Zone 6 | 28,53 | 12,9 | 15,26 | 17,54 | 24,61 | 22,84 | | | | |
| Zone 7 | 22,93 | 11,29 | 11,18 | 16,2 | 24,62 | 20,46 | 23,62 | | | |
| Zone 8 | 24,59 | 9,75 | 11,56 | 12,33 | 15,48 | 18,48 | 21,7 | 17,29 | | |
| Alpine zone | 22,7 | 5,78 | 6,97 | 7,23 | 10,01 | 11,31 | 15,51 | 12,26 | 18,98 | |

SIMPER(1.7%)

| | Within Zone Similarity (%) | Zone 1 | Zone 2 | Zone 3 | Zone 4 | Zone 5 | Zone 6 | Zone 7 | Zone 8 | Alpine zone |
|---|---|---|---|---|---|---|---|---|---|---|
| | | Between Zone Similarity (%) | | | | | | | | |
| Zone 1 | 26,58 | | | | | | | | | |
| Zone 2 | 26,76 | 21,17 | | | | | | | | |
| Zone 3 | 31,00 | 24,78 | 23,12 | | | | | | | |
| Zone 4 | 31,12 | 18,54 | 22,09 | 23,16 | | | | | | |
| Zone 5 | 30,25 | 15,65 | 16,83 | 18,99 | 25,22 | | | | | |
| Zone 6 | 30,51 | 15,45 | 19,30 | 19,76 | 28,20 | 26,54 | | | | |
| Zone 7 | 24,63 | 13,79 | 15,01 | 17,82 | 27,61 | 23,90 | 26,57 | | | |
| Zone 8 | 27,83 | 12,03 | 14,59 | 14,94 | 20,01 | 22,94 | 24,43 | 21,48 | | |
| Alpine zone | 27,85 | 7,82 | 10,58 | 9,13 | 14,20 | 16,96 | 18,58 | 16,21 | 23,34 | |

SIMPER(3%)

| | Within Zone Similarity (%) | Zone 1 | Zone 2 | Zone 3 | Zone 4 | Zone 5 | Zone 6 | Zone 7 | Zone 8 | Alpine zone |
|---|---|---|---|---|---|---|---|---|---|---|
| | | Between Zone Similarity (%) | | | | | | | | |
| Zone 1 | 28,65 | | | | | | | | | |
| Zone 2 | 29,34 | 24,24 | | | | | | | | |
| Zone 3 | 33,58 | 26,57 | 24,54 | | | | | | | |
| Zone 4 | 32,51 | 22,1 | 25,27 | 25,83 | | | | | | |
| Zone 5 | 32,95 | 21,05 | 21,4 | 23,21 | 28,28 | | | | | |
| Zone 6 | 21,12 | 19,95 | 21,83 | 22,79 | 30,33 | 29,28 | | | | |
| Zone 7 | 26,49 | 17,75 | 17,75 | 21,25 | 30,69 | 25,18 | 28,8 | | | |
| Zone 8 | 32,74 | 19,43 | 20,89 | 20,63 | 25,54 | 26,51 | 27,94 | 24,72 | | |
| Alpine zone | 42,98 | 19,01 | 22 | 18,42 | 22,23 | 23,19 | 25,82 | 22,07 | 31,1 | |

SIMPER(4%)

| | Within Zone Similarity (%) | Zone 1 | Zone 2 | Zone 3 | Zone 4 | Zone 5 | Zone 6 | Zone 7 | Zone 8 | Alpine zone |
|---|---|---|---|---|---|---|---|---|---|---|
| | | Between Zone Similarity (%) | | | | | | | | |
| Zone 1 | 31,14 | | | | | | | | | |
| Zone 2 | 31,54 | 26,99 | | | | | | | | |
| Zone 3 | 36,58 | 28,86 | 26,95 | | | | | | | |
| Zone 4 | 35,64 | 24,58 | 27,98 | 26,58 | | | | | | |
| Zone 5 | 34,36 | 24,07 | 26,03 | 24,9 | 30,76 | | | | | |
| Zone 6 | 34,78 | 22,83 | 25,2 | 24,48 | 33,88 | 32,12 | | | | |
| Zone 7 | 29,92 | 20,92 | 21,8 | 23,14 | 33,29 | 27,78 | 31,91 | | | |
| Zone 8 | 36,37 | 23,26 | 25,28 | 22,51 | 31,01 | 30,97 | 31,34 | 29,95 | | |
| Alpine zone | 50,63 | 23,93 | 27,77 | 20,88 | 29,72 | 30,16 | 30,24 | 28,48 | 36,41 | |

SIMPER(5%)

| | Within Zone Similarity (%) | Zone 1 | Zone 2 | Zone 3 | Zone 4 | Zone 5 | Zone 6 | Zone 7 | Zone 8 | Alpine zone |
|---|---|---|---|---|---|---|---|---|---|---|
| | | Between Zone Similarity (%) | | | | | | | | |
| Zone 1 | 35,24 | | | | | | | | | |
| Zone 2 | 39,81 | 32,44 | | | | | | | | |
| Zone 3 | 38,77 | 30,97 | 30,71 | | | | | | | |
| Zone 4 | 39,31 | 28,82 | 34,37 | 28,22 | | | | | | |
| Zone 5 | 37,94 | 27,45 | 33,14 | 27,08 | 36 | | | | | |
| Zone 6 | 41,08 | 28,29 | 33,91 | 28,38 | 39,25 | 36,16 | | | | |
| Zone 7 | 35,39 | 27,73 | 31,71 | 28,12 | 39,45 | 33,31 | 37,9 | | | |
| Zone 8 | 42,79 | 28,78 | 34,95 | 26,97 | 37,83 | 36,36 | 37,91 | 35,46 | | |
| Alpine zone | 62,06 | 31,66 | 40,94 | 28,65 | 39,99 | 38,97 | 39,78 | 38,92 | 45,42 | |

### Supplementary Table S4. Distinctness of scuttle fly communities across time periods.

*Shown are results of ANOSIM analyses of species turnover across time-periods for all zones (top) and northern (IV-alpine) zones only (bottom).*

| ANOSIM | R=0.269, P=0.001 | | | |
|---|---|---|---|---|
| | Late Spring | Midsummer | Late Summer | Offseason |
| Late Spring | | 0,091 | 0,350 | 0,390 |
| Midsummer | 0,018 | | 0,262 | 0,233 |
| Late Summer | 0,001 | 0,001 | | 0,339 |
| Offseason | 0,001 | 0,002 | 0,001 | |

| ANOSIM | R=0.588, P=0.001) | | | |
|---|---|---|---|---|
| | Late Spring | Midsummer | Late Summer | Offseason |
| Late Spring | | 0.104 | 0,882 | 0,728 |
| Midsummer | 0,125 | | 0,712 | 0,383 |
| Late Summer | 0,001 | 0,001 | | 0,826 |
| Offseason | 0,001 | 0,003 | 0,001 | |

### Supplementary Table S5. Similarity of scuttle fly communities across time periods.

*Shown are results of SIMPER analyses of species turnover across time-periods for all zones (top) and northern (IV-alpine) zones only (bottom).*

| SIMPER | Within Season | Between Season Similarity (%) | | | |
|---|---|---|---|---|---|
| | Similarity (%) | Late Spring | Midsummer | Late Summer | Offseason |
| Late Spring | 26,52 | | | | |
| Midsummer | 23,57 | 22,12 | | | |
| Late Summer | 20,61 | 14,45 | 16,40 | | |
| Offseason | 24,23 | 16,58 | 18,59 | 13,12 | |

| SIMPER | Within Season | Between Season Similarity (%) | | | |
|---|---|---|---|---|---|
| | Similarity (%) | Late Spring | Midsummer | Late Summer | Offseason |
| Late Spring | 42,93 | | | | |
| Midsummer | 32,81 | 31,55 | | | |
| Late Summer | 29,30 | 10,62 | 16,25 | | |
| Offseason | 43,58 | 29,03 | 26,45 | 13,87 | |

### Supplementary Table S6. Mantel tests relating pairwise community dissimilarity to pairwise differences in space, time or both.

*To examine the effect of the species delimitation criteria used, we computed pairwise Jaccard community dissimilarity values using both individual haplotypes (dataset "Haplotypes") and species defined by 1.7% sequence divergence (dataset "Species"). Total dissimilarity (Total Betadiversity) was resolved into its Turnover and Nestedness components. Each component was then related to the spatial difference (i.e., the geographical distance between sites), the temporal difference (i.e., temporal distance in weeks between samples) and the spatio-temporal differences (i.e., geographical distance and temporal distance, including their joint effect) between sample pairs. Columns* r *denote Pearson moment product correlations, whereas* p-*values are based on Mantel tests.*

| Dataset | Distances | Turnover | | Nestedness | | Total Betadiversity | |
|---|---|---|---|---|---|---|---|
| | | *r* | *p* | *r* | *p* | *r* | *p* |

| | | | | | | | |
|---|---|---|---|---|---|---|---|
| Species | Spatial | **0.75** | **0.001** | -0.33 | 1 | **0.65** | **0.001** |
| | Temporal | -0.03 | 0.5 | 0.53 | 0.21 | 0.58 | 0.125 |
| | SpatioTemporal – Late spring | **0.57** | **0.001** | -0.19 | 0.997 | **0.64** | **0.001** |
| | SpatioTemporal – Midsummer | **0.64** | **0.001** | -0.31 | 1 | **0.61** | **0.001** |
| | SpatioTemporal – Late Summer | **0.74** | **0.001** | -0.41 | 1 | **0.73** | **0.001** |
| | SpatioTemporal – Offseason | **0.25** | **0.001** | -0.25 | 1 | **0.43** | **0.001** |
| Haplotypes | Spatial | **0.79** | **0.001** | -0.37 | 1 | **0.69** | **0.001** |
| | Temporal | -0.73 | 0.92 | 0.93 | 0.083 | **0.69** | **0.041** |
| | SpatioTemporal – Late spring | **0.64** | **0.001** | -0.34 | 1 | **0.60** | **0.001** |
| | SpatioTemporal – Midsummer | **0.63** | **0.001** | -0.24 | 1 | **0.62** | **0.001** |
| | SpatioTemporal – Late Summer | **0.72** | **0.001** | -0.43 | 1 | **0.71** | **0.001** |
| | SpatioTemporal – Offseason | **0.14** | **0.001** | -0.17 | 1 | **0.42** | **0.001** |
